# Genomic and Transcriptomic Landscapes of MEN1-Wild-Type Low-Grade Metastatic Pancreatic NETs Uncover Key Oncogenic Drivers and Targetable Pathways

**DOI:** 10.64898/2025.12.07.692731

**Authors:** Md Hafiz Uddin, Zaid Mahdi, Irfana Muqbil, Brendon R. Herring, Bart Rose, Husain Y. Khan, Yiwei Li, Amro Aboukameel, Sahar F. Bannoura, Hugo Jimenez, Allan Johansen, Mohammad Najeeb Al-Hallak, Ibrahim Azar, Amr Mohamed, Tarik Hadid, Nitin Vaishampayan, Yang Shi, Yin Wan, Vy Ong, Gregory Dyson, Rafic Beydoun, Miguel Tobon, Eliza W. Beal, Herbert Chen, Anthony F. Shields, Philip A. Philip, Jennifer Beebe-Dimmer, Ramzi M. Mohammad, Boris C. Pasche, Bassel F. El-Rayes, Asfar S. Azmi

**Affiliations:** Karmanos Cancer Institute, Detroit, MI; Emory Winship Cancer Institute, Atlanta, GA; Lawrence Technological University, Southfield, MI; O’Neal Comprehensive Cancer Center, University of Alabama at Birmingham, Birmingham, AL; Department of Cancer Biology, Wake Forest University, Winston-Salem, NC; Trinity Health Oakland, Pontiac, MI; UH Seidman Cancer Center, University Hospitals, Case Western Reserve University, Cleveland, OH; Department of Pathology, Wayne State University School of Medicine, Wayne State University, Detroit, MI; Henry Ford Health System, Detroit, MI

**Keywords:** Pancreatic neuroendocrine tumors, genomic and transcriptomic characterization, low grade, metastasis, mutation, therapeutic candidates

## Abstract

Sporadic pancreatic neuroendocrine tumors (pNETs) with wild type MEN1 represent a major yet largely ignored subset whose biology and metastatic potential remain poorly understood. Because metastasis can occur despite low histologic grade and modest mutational burden, we hypothesized that metastatic competence in MEN1-wild-type pNETs reflects quantitative reinforcement of shared oncogenic pathways rather than distinct mutational processes. We profiled 75 primary low-grade pNETs by whole-exome and RNA sequencing, including 25 percent with lymph node and/or liver metastasis, and integrated genomic and transcriptomic data to connect pathway lesions with expression state. Metastatic tumors showed a slight increase in mutation frequency but conserved base-substitution spectra relative to non-metastatic cases, and adverse clinicopathologic features were enriched in Grade 2 disease. Aggregating alterations to pathways revealed broad convergence on canonical networks, with transcriptomic analyses demonstrating cohort-wide enrichment of Calcium, WNT, and KRAS/PI3K-AKT programs in metastasis. Intersection of significantly mutated genes with differentially expressed genes identified a focused 29-gene overlap, including RYR1 and ZNF273, that marks these convergent axes and distinguishes metastatic from non-metastatic tumors. Gene set enrichment confirmed preferential activation of Calcium, WNT, and PI3K-AKT signaling in metastatic tumors, consistent with a network-intensity model of progression. Finally, upstream-regulator analysis (iPathwayGuide) and gene-centric perturbation mapping (Gene2Drug) nominated candidate targeted and repurposable agents predicted to reverse the metastatic expression phenotype and flagged drugs unlikely to provide benefit, yielding a prioritized, testable therapeutic shortlist which includes fasudil and spaglumic acid. Convergent, domain-specific mutational patterns in highly mutated genes such as ZNF273 and CLCA1 define a molecular signature that could stratify metastatic risk in low-grade pNETs. Collectively, our data reframe metastasis in MEN1-wild-type low-grade pNETs as a property of pathway state rather than mutation quantity and provide a translational blueprint for biomarker-guided therapy development focused on Calcium, WNT, and KRAS/PI3K hubs.

## Introduction

Neuroendocrine tumors (NETs) arise from the hormone-producing cells of various organs throughout the body and include pancreatic NETs (pNETs), medullary thyroid cancer, gastrointestinal (GI) NETs, bronchopulmonary NETs, and paraganglioma/pheochromocytoma^1^. Over 50% of patients with pancreatic NETs develop isolated hepatic metastases^1^. According to the American Cancer Society’s estimate, in the United States over 4,000 individuals were diagnosed with pNETs in 2020 (ACS Statistics)^2^. Overall survival for pNETs can be relatively long compared to other cancers, leading to a higher number of cases of this disease at any given time^3,4^. Pancreatic neuroendocrine tumors (pNETs) are heterogeneous neoplasms with rising incidence and substantial clinical variability, ranging from indolent lesions to tumors that metastasize despite low histologic grade, thus the management of pNETs remains clinically challenging^2^. Improvements in the identification of actionable oncogenic drivers or systemic treatments for this growing patient population have been at best modest in recent decades. Therefore, pNETs remain a significant unmet clinical problem and in urgent need of newer biomarkers as well as more effective therapeutics.

pNETs can result from heritable genetic or somatic non-familial mutations^5,6^. Earlier, large scale genomic studies revealed that loss of Multiple Endocrine Neoplasia Type 1 (MEN1), Death-Domain-Associated protein (DAXX), and α Thalassemia/mental Retardation Syndrome X-linked (ATRX) genes as the major genetic aberrations in pNETs^7^. Additionally, the hyperactivation of PI3K-Akt-mTOR through loss of tumor suppressor PTEN has been well documented as a main driver in pNETs^8,9^. Molecular studies over the past decade have defined recurrent alterations in MEN1, DAXX, and ATRX, and frequent perturbations of PI3K-mTOR signaling, establishing a canonical framework for tumorigenesis and motivating targeted therapy trials^7,10–13^. These genomic advances translated into clinically meaningful gains with agents such as everolimus^14^ and sunitinib, which prolong progression-free survival in advanced disease^12^, yet durable control and robust biomarkers remain limited ^11,15,16^. More recent multi-omic and population-level analyses underscore that tumor evolution and outcome are not fully explained by the presence or absence of MEN1/DAXX/ATRX lesions alone, and that broader network-level perturbations likely shape malignant potential and therapeutic response^17–19^. Within this landscape, MEN1-wild type disease constitutes a major and under characterized subset of sporadic pNETs in which the determinants of metastatic behavior and drug sensitivity remain unclear.

Despite progress in cataloging mutations, key biological and clinical questions persist^20^. Much of the field’s mechanistic understanding derives from cohorts enriched for MEN1-mutant tumors or from studies that emphasize single-gene drivers rather than pathway state, leaving MEN1-wild type pNETs relatively unresolved^18,21,22^. While PI3K-mTOR axis activity is common, it incompletely predicts metastatic risk or treatment benefit, implying that additional signaling circuits and microenvironmental interactions modulate aggressiveness^17,23^. The tumor mutational burden in pNETs is generally low to moderate and does not consistently stratify outcomes, raising the possibility that metastatic competence may arise from quantitative reinforcement of shared oncogenic pathways rather than qualitative shifts in mutational processes^24^.

To address these gaps, we assembled and profiled a rigorously curated cohort of MEN1-wild type low-grade pNETs using matched whole-exome and RNA sequencing and integrated the resulting data to connect genomic lesions with transcriptional state. Our guiding hypothesis was that metastatic behavior in histologically low-grade tumors reflects convergence on a limited set of oncogenic pathways that are quantifiably more active in metastasis and that these pathway states can be mapped to pharmacologic space using complementary in silico approaches. We therefore contrasted non-metastatic and metastatic cases, aggregated mutations to pathway-level signatures, and asked which axes show concordant genomic/transcriptomic deregulation and cohort-wide enrichment by gene set analysis.

Our integrated analysis supports a revised view of aggressiveness in MEN1-wild type pNETs. We show that metastatic tumors exhibit only modest increases in mutation frequency relative to non-metastatic counterparts and conserve base-substitution spectra, yet display stronger engagement of convergent signaling programs, notably Calcium signaling, WNT, and KRAS/PI3K-related pathways, with a small set of genes showing overlap between somatic alteration and differential expression. Our framework integrates pathway-level mutation maps with transcriptomic enrichment and drug-prediction analyses to move beyond catalogs toward actionable network states. Current study provides a translational bridge by applying iPathwayGuide and Gene2Drug to nominate candidate therapeutics that are predicted to reverse the metastatic expression phenotype and to identify agents unlikely to help in this context. The methodology is supported by prior validations of these tools for mechanistic inference and drug prioritization, and by reports implicating RYR1 dysregulation and related calcium handling machinery in tumor biology across solid cancers^25–28^. Together, this study reframes metastatic propensity as a property of network intensity and transcriptional reinforcement rather than new mutational categories, and it outlines how integrated genomics and transcriptomics can prioritize therapies for a clinically important yet understudied subset of pNETs.

## Methods

### Collection, processing and sequencing of patient samples

Under an IRB approved protocol (STUDY00001739), a total of 104 Formalin-Fixed, Paraffin-Embedded (FFPE) tissues (98 low grade pancreatic neuroendocrine tumor (pNET) and six normal pancreatic) from the Emory University tissue biobank were sectioned and the genomic DNA was extracted using QIAamp DNA FFPE Tissue Kit (Qiagen, Germany). Sex was considered as a biological variable and average age of the patients was 55.83 years. Among the 98 pNET tissues, eight samples did not have information related to grade and stage were therefore excluded. Next, eight samples failed QC and were excluded. Finally, seven samples are not classified as Grade 1 or 2 therefore excluded (Figure 1A shows tissue selection schema).

**Figure 1.**
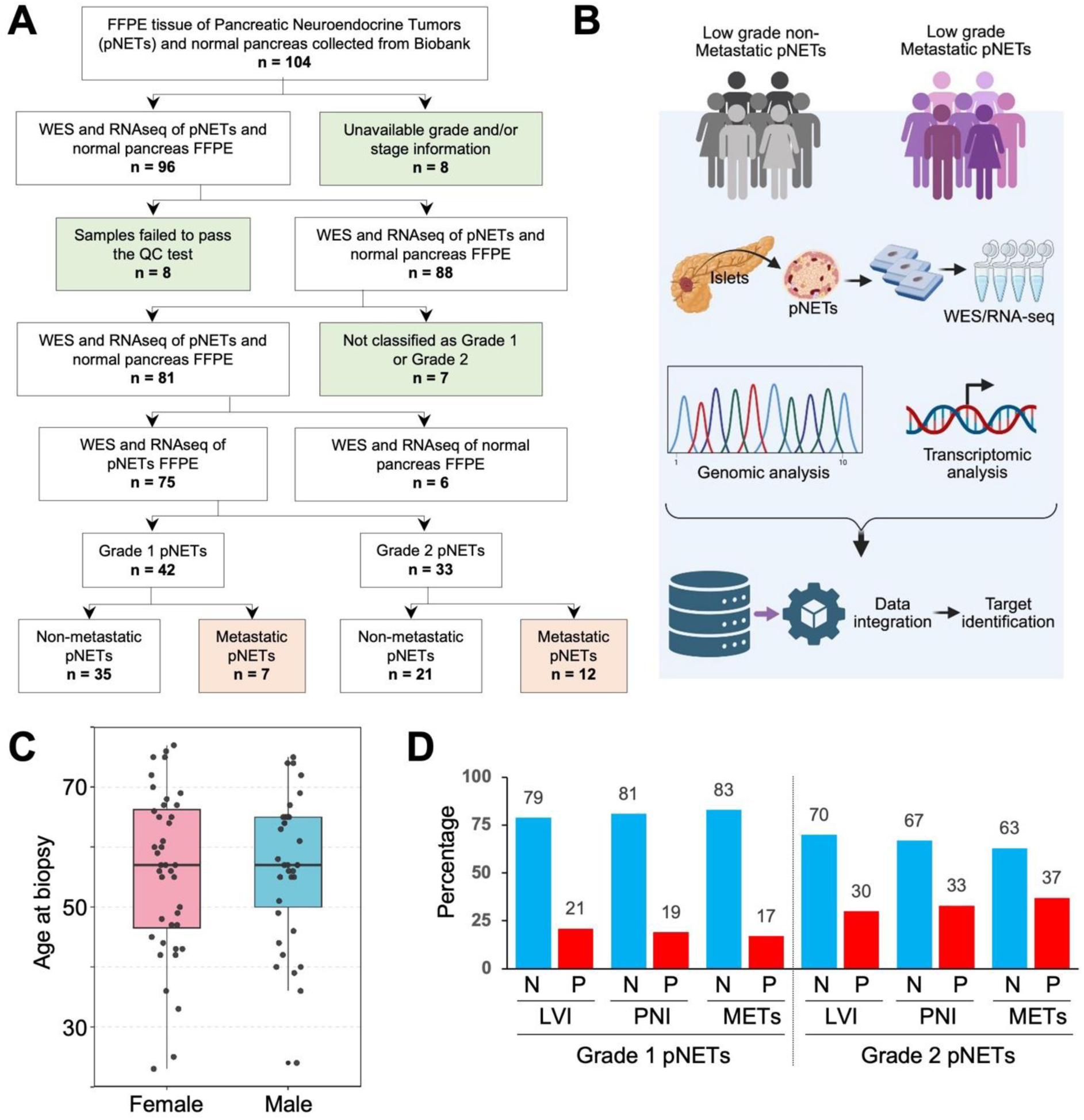
Overview of patient inclusion and subclassification FFPE tissues. **A.** Flowchart of patient inclusion. From low grade pNETs patient cohort, FFPE biopsy tissues from 104 distinct patients were selected. Total pool of samples was subject to whole exome sequencing (WES) and RNA sequencing (RNA-seq). The filtering criteria that led to sample exclusion are highlighted in light green. The final set of selected samples is depicted in white boxes. Metastatic samples are highlighted in light pink. **B.** Subclassification of pNETs patients and samples processing schema. The 75 primary pNETs were subclassified into two major categories based on their metastatic outcome. **C.** Age distribution stratified by gender of the pNET cohort. The variable’s distribution and median were visualized with data points clustered on the boxplot. **D.** Percentage of lympho-vascular invasion (LVI), peri-neural invasion (PNI) and metastasis (METs) in Grade 1 and Grade 2 pNET patients. Exact percentage shown on top of each bar. N, negative (blue); P, positive (red). Credit: Figure 1B was Created with BioRender (https://biorender.com/).

### Collection of pathological records

Primary tumor characteristics of the pNET patients were obtained from the histo- and cytopathology registry at Emory University. Data collected included patient gender, WHO-defined tumor differentiation grade (Grade 1 or Grade 2), proliferation index (Ki-67), and evidence of invasion or metastasis. For each patient, information was extracted from the pathological reports of the primary and/or metastatic lesions. When multiple reports were available, the record closest in date but always preceding the biopsy was considered.

### Whole exome sequencing

For whole exome sequencing (WES), DNA samples from pNET and normal tissues were fragmented by using sonication and subjected to library construction in blinded fashion. Exome capture was performed using the SureSelect Human All Exon V6 Kit (Agilent Technologies) following the vendor’s protocol. Sequencing was performed using the Illumina Hiseq X Ten with 150-bp paired-end reads for a total of 40 million reads. Prior to alignment, the low-quality reads were removed. Reads were aligned to the reference genome using Burrows-Wheeler Alignment (BWA). Picard tools were utilized to identify and remove duplicate reads from the BAM files generated by BMA. The resulting BAM files were indexed and sorted by samtools.

### Mutational signature analysis

The Genome Analysis Toolkit (GATK) was utilized to identify the genomic variants in the WES data. Somatic variant calls were generated with GATK Mutect2, which assigned an accurate confidence score to each putative mutation call and evaluated new potential variants. Biological functional annotations for the identified somatic variants were performed using Annovar. Somatic copy number variants (CNVs) were detected using Facets. The R Bioconductor package, Maftools were utilized for summarizing, analyzing and visualizing. Mutation Annotation Format (MAF) files generated from GATK, which created the oncoplots, transition and transversion plots, and lollipop mutation diagrams to visualize the somatic mutations in pNET patients.

### RNA-sequencing

Total RNA was extracted from pNET and normal tissues using the miRNeasy mini kit from Qiagen. The primary pNETs and pancreatic normal FFPE (Formalin-Fixed, Paraffin-Embedded) tissues were sectioned, and the total RNA was extracted following manufacturer’s instruction (Qiagen, Germany). Total RNA-sequencing was conducted by LC Sciences. For total RNA sequencing, RNA integrity was checked with Agilent Technologies 2100 Bioanalyzer to ensure RIN >7 cutoff value. All the samples passed the quality control except four pNET samples. Approximately 10 µg of total RNA was used to isolate poly(A) mRNA employing poly-T oligo–conjugated magnetic beads (Invitrogen). The captured poly(A)+ or poly(A)- RNA fractions were fragmented into short fragments using divalent cations under elevated temperature. The resulting RNA fragments were then reverse transcribed to generate complementary DNA (cDNA) libraries according to the manufacturer’s protocol (Illumina, San Diego, CA, USA). The sequence library was prepared following Illumina’s TruSeq-stranded-total-RNA-sample preparation protocol. Quality control analysis and quantification of the sequencing library were performed using Agilent Technologies 2100 Bioanalyzer High Sensitivity DNA Chip. Paired-end sequencing was performed on Illumina’s NovaSeq 6000 sequencing system for a total of 40 million reads per sample. The reads in Fastq format were mapped to the human genome using HIAST2. Expression levels of mRNAs were quantified using StringTie^29^ based on fragments per kilobase of transcript per million mapped reads (FPKM), and the expression of transcripts generated from the same gene were aggregated to obtain the expression level of the gene. Differentially expressed genes were determined using the R Bioconductor package, applying the thresholds of absolute values of log2 (fold change) > 1 and a false discovery rate (FDR) < 0.05 for statistical significance.

### Inventory of clinically actionable putative therapeutic targets against somatic alterations

We prioritized candidate therapies by integrating pathway-level upstream regulator inference (iPathwayGuide, Advaita Bio; AKB v18.1, 2025 release) with gene-centric drug perturbation profiling (Gene2Drug, accessed 2025) using the same tumor set. Differentially expressed genes (DEGs) were derived from metastatic versus non-metastatic low-grade pNETs with gene symbols harmonized to HGNC. In iPathwayGuide, DEGs were mapped onto the Advaita Knowledge Base and analyzed via impact analysis, which combines (i) over-representation statistics and (ii) perturbation of pathway topology to model directionality across signaling cascades. In parallel, Gene2Drug queried the subset of “overlapped” genes (significantly mutated and differentially expressed in the same tumors) plus pathway-adjacent DEGs against LINCS/Connectivity-Map derived perturbational signatures across multiple annotation spaces (cancer gene perturbations, Gene Ontology, transcription factor target sets etc). For each compound, an enrichment score was computed over rank-ordered drug-response profiles.

### Statistical analysis

All analyses were performed on primary low-grade pNETs with matched WES and RNA-seq. For RNA-Seq, differential expressed genes were assessed with the R Bioconductor package edgeR with the thresholds of absolute value of log2 (fold change) > 1 and FDR < 0.05. For WES data, tumor mutational burden (TMB) was computed as somatic single-nucleotide variants and short indels per callable megabase in the exome; base-substitution spectra were summarized in the six canonical SNV classes. Differential mutated genes between the pNET and normal samples were identified using the Fisher’s exact test with a p-value < 0.05 and the difference in mutation frequencies between the two groups greater than 20%. The R Bioconductor package clusterProfiler was used for functional enrichment analysis, where the over-representation tests were used for KEGG (Kyoto Encyclopedia of Genes and Genomes) pathway and Gene Ontology (GO) categories used and gene set enrichment analysis (GSEA) was performed for the Molecular Signatures Database (MSigDB) Hallmark gene set collections. Significantly enriched pathways, GOs and gene sets were identified using FDR< 0.05.

## Results

### Clinicopathologic stratification of low-grade pNETs for integrated genomic and transcriptomic analysis

From the institutional cohort of low-grade pNETs, we identified 104 patients with available FFPE biopsy material and subjected all corresponding specimens to whole-exome sequencing (WES) and RNA sequencing (RNA-seq) (**Figure 1A**). Samples were sequentially filtered to ensure clinicopathologic and molecular data quality. Twenty-three cases were removed because they failed prespecified QC metrics for nucleic acid yield or sequencing performance, lacked reliable information on grade and/or stage, or could not be unequivocally classified as WHO Grade 1 or Grade 2. After these exclusions, low-grade pNETs with high-quality genomic and transcriptomic data were retained for downstream analyses. Metastatic cases within this refined cohort are highlighted in **Figure 1A** (light pink boxes).

To investigate molecular determinants of metastatic behavior, the primary tumors were subsequently stratified into two major categories according to their metastatic outcome (**Figure 1B**) including low-grade non-metastatic and metastatic pNETs. Both groups underwent a unified workflow, including DNA and RNA extraction from FFPE blocks, WES and RNA-seq, and integrated genomic-transcriptomic analysis for candidate target identification. Among the primary tumors, 7 Grade-1 and 12 Grade-2 pNETs were with metastatic status (**Figure 1A**), providing a clinically annotated set of cases for comparative analyses with non-metastatic counterparts.

The demographic and pathological characteristics of the sequenced cohort are summarized in **Figures 1C-1D** and **Table 1**. Age at diagnosis displayed a wide distribution in both sexes, with overlapping ranges and comparable median ages for male and female patients (**Figure 1C**), indicating that metastatic propensity in this cohort is not driven by major age or sex imbalances. In contrast, adverse histopathologic features were enriched among Grade 2 tumors. Grade 2 tumors exhibited substantially higher rates of lympho-vascular invasion (LVI), peri-neural invasion (PNI), and metastasis (METs) compared to Grade 1 (30-37% positive vs. 17-21% in Grade 1; **Figure 1D**). Collectively, these data define a well-curated set of primary low-grade pNETs with linked clinical, histologic, and multi-omic profiles, and underscore the association between metastatic potential and genomic/transcriptomic characteristics.

**Table 1:**
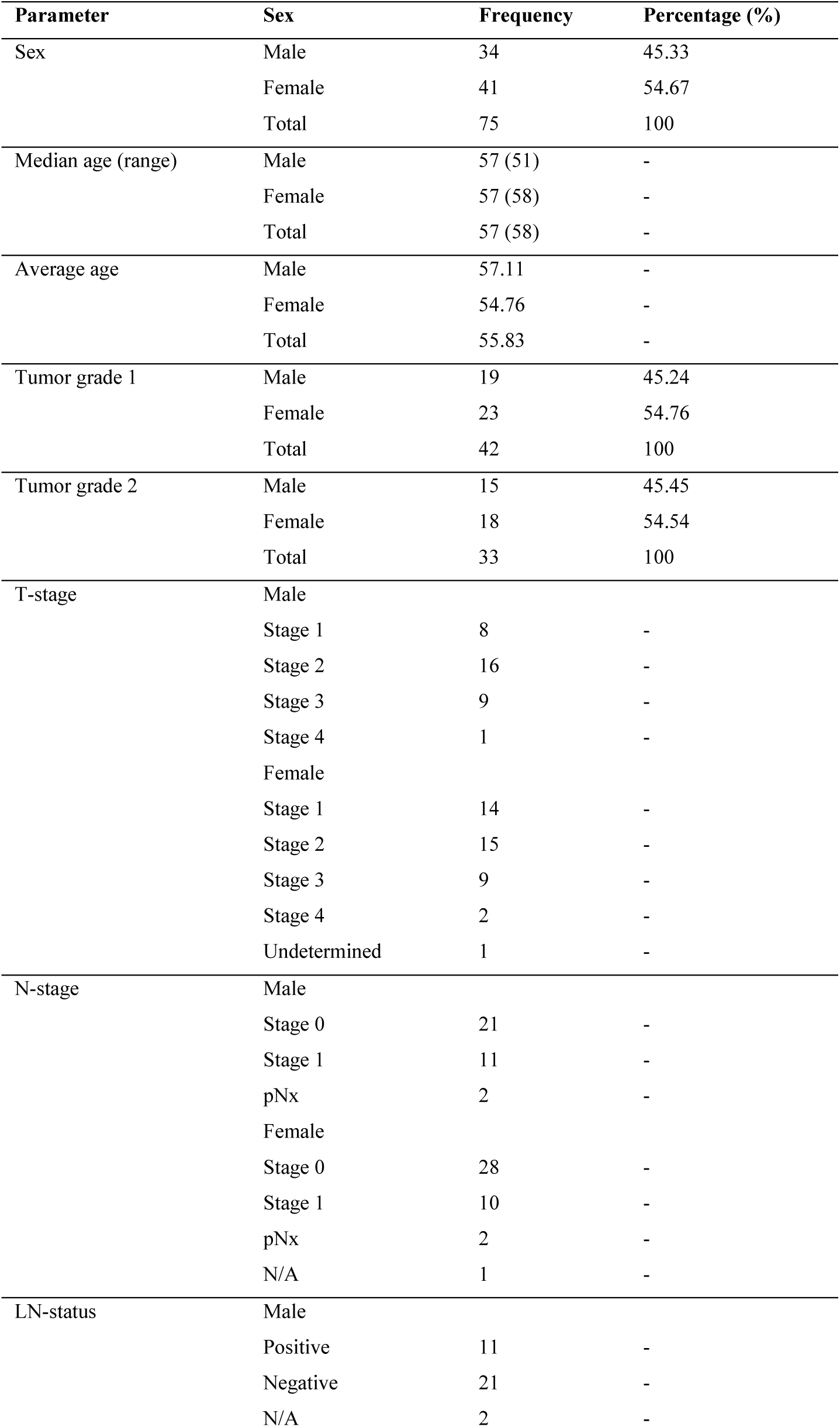
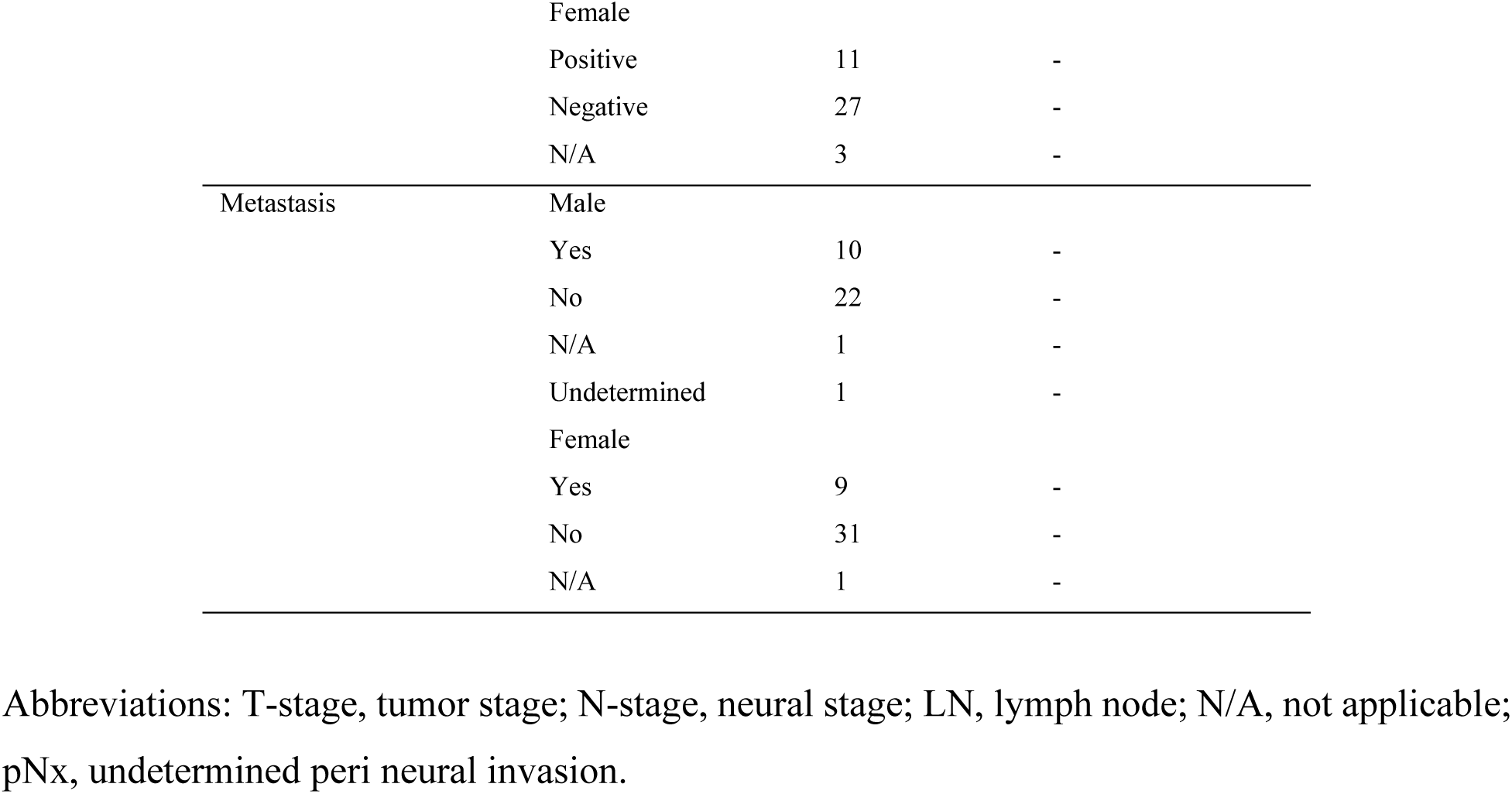
Clinical and Demographic Characteristics of Low-Grade pNET Patients.

| Parameter | Sex | Frequency | Percentage (%) |
| --- | --- | --- | --- |
| Sex | Male | 34 | 45.33 |
|  | Female | 41 | 54.67 |
|  | Total | 75 | 100 |
| Median age (range) | Male | 57 (51) | - |
|  | Female | 57 (58) | - |
|  | Total | 57 (58) | - |
| Average age | Male | 57.11 | - |
|  | Female | 54.76 | - |
|  | Total | 55.83 | - |
| Tumor grade 1 | Male | 19 | 45.24 |
|  | Female | 23 | 54.76 |
|  | Total | 42 | 100 |
| Tumor grade 2 | Male | 15 | 45.45 |
|  | Female | 18 | 54.54 |
|  | Total | 33 | 100 |
| T-stage | Male |  |  |
|  | Stage 1 | 8 | - |
|  | Stage 2 | 16 | - |
|  | Stage 3 | 9 | - |
|  | Stage 4 | 1 | - |
|  | Female |  |  |
|  | Stage 1 | 14 | - |
|  | Stage 2 | 15 | - |
|  | Stage 3 | 9 | - |
|  | Stage 4 | 2 | - |
|  | Undetermined | 1 | - |
| N-stage | Male |  |  |
|  | Stage 0 | 21 | - |
|  | Stage 1 | 11 | - |
|  | pNx | 2 | - |
|  | Female |  |  |
|  | Stage 0 | 28 | - |
|  | Stage 1 | 10 | - |
|  | pNx | 2 | - |
|  | N/A | 1 | - |
| LN-status | Male |  |  |
|  | Positive | 11 | - |
|  | Negative | 21 | - |
|  | N/A | 2 | - |
|  | Female |  |  |
|  | Positive | 11 | - |
|  | Negative | 27 | - |
|  | N/A | 3 | - |
| Metastasis | Male |  |  |
|  | Yes | 10 | - |
|  | No | 22 | - |
|  | N/A | 1 | - |
|  | Undetermined | 1 | - |
|  | Female |  |  |
|  | Yes | 9 | - |
|  | No | 31 | - |
|  | N/A | 1 | - |

Abbreviations: T-stage, tumor stage; N-stage, neural stage; LN, lymph node; N/A, not applicable; pNx, undetermined peri neural invasion.

### Comprehensive landscape of somatic mutations in low-grade pNETs reveals conserved mutational signatures across metastatic states

The global mutational landscape of primary low-grade pNETs was next examined by ordering tumors according to metastatic outcome and histologic grade (**Figure 2**). Across the sequenced tumors, the overall tumor mutational burden (TMB) in exons including upstream, downstream and non-coding sequences was moderate, with most cases harboring relatively low number of mutations per megabase (Mb). A small subset of outliers exhibiting markedly elevated TMB (**Figure 2A**). A similar pattern was observed when analysis was restricted to coding regions, where exonic TMB paralleled the distribution of overall TMB (**Figure 2C**). Hypermutated tumors were observed in both metastatic and non-metastatic groups, however, average mutation per Mb were slightly higher in pNETs with metastatic status.

**Figure 2.**
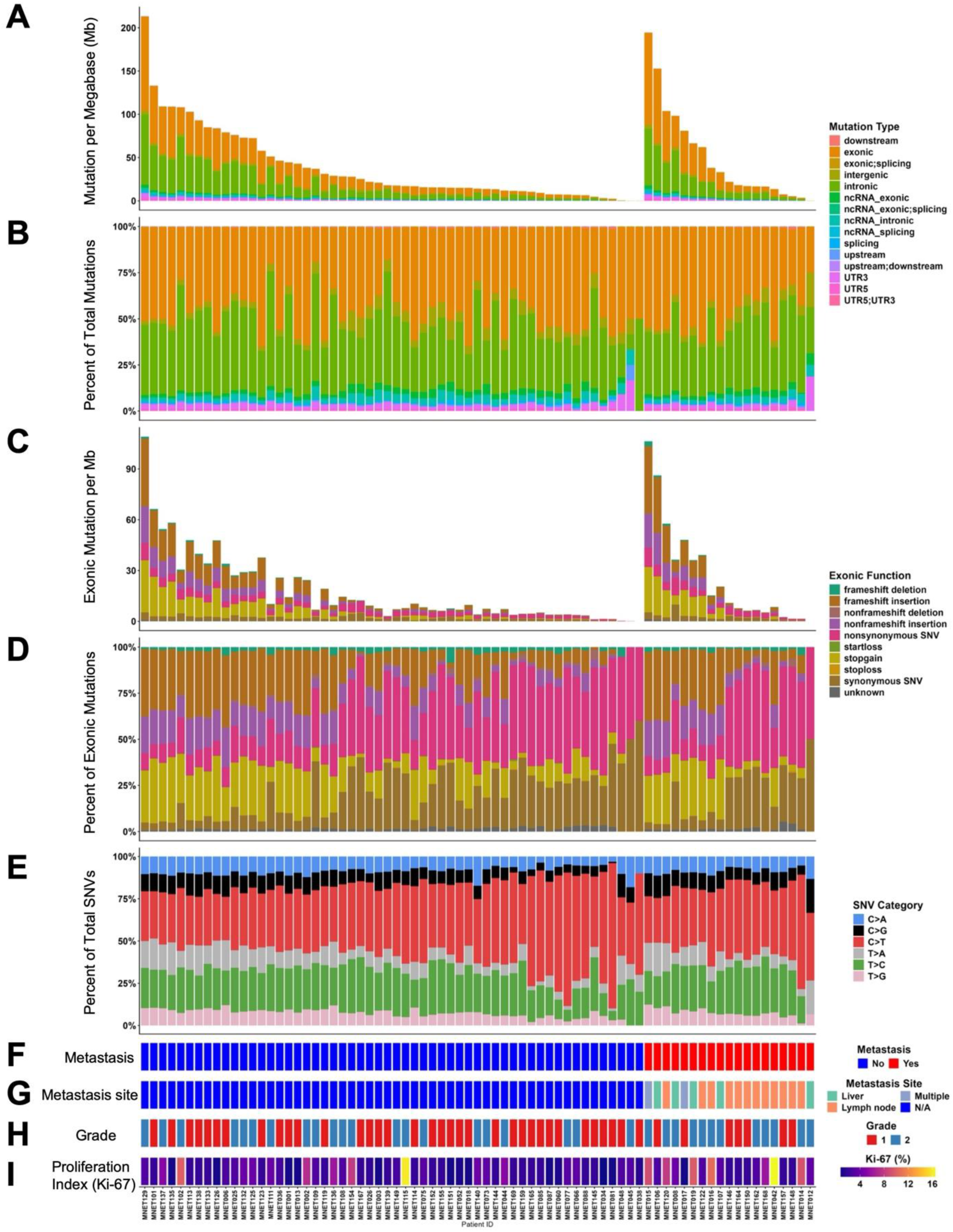
Landscape of large-scale genomic alterations detected in pNETs, ordered by metastatic potential and differentiation grade. **A.** Overview of genomic characteristics of pNETs, tumor mutational burden (TMB) ordered by metastatic potential and differentiation grade.For each pNET (n = 75), a number of genomic mutations per megabase (Mb) is shown. B. Percentage of mutations for all mutation types in all pNET samples. **C.** Overview of exonic TMB ordered by metastatic potential and differentiation grade. For each pNET (n = 75), a number of exonic Mb is shown. **D.** Percentage of exonic mutations for all mutation types in all pNET samples. **E.** Relative frequency (percentage) of pyrimidine point mutation as the total single nucleotide variants (SNVs) with six categories for all pNET samples. **F.** Categorization of primary tumors based on the patient’s metastatic status. Metastatic patient samples are depicted in red and non-metastatic patient samples are depicted in blue. **G.** Categorization of metastasis based on their sites. Liver metastasis depicted in green, lymph node metastasis depicted in orange, metastasis in multiple sites depicted in light purple and non-metastatic samples marked in blue. **H.** Differentiation grade of the pNETs; grade 1 (G1) samples are shown in red and grade 1 (G1) samples are shown in blue. **I.** Proliferation index (Ki-67) from the pathological record where scores range 0-16 are represented as a gradient between dark purple as the lowest score to yellow as the highest score.

Inspection of mutation annotations showed that the majority of variants resided in non-coding regions, predominantly intronic and intergenic, whereas exonic variants accounted for a moderate fraction of the total mutational burden (**Figure 2B**). Within coding regions, nonsynonymous single-nucleotide variants (SNVs) and frameshift or non-frameshift insertions/deletions represented the dominant functional classes (**Figure 2D**). Synonymous substitutions and start/stop-altering events were comparatively infrequent, and the relative proportions of these exonic functional classes were broadly similar across tumors. Analysis of base-substitution spectra revealed a characteristic pyrimidine mutation pattern dominated by C>T transitions, with additional contributions from T>C and C>A changes (**Figure 2E**). This six-class SNV profile was highly consistent across the cohort and did not show obvious qualitative shifts between metastatic and non-metastatic tumors, suggesting that the operative mutational processes are shared across clinical subgroups.

To relate these genomic features to clinicopathologic parameters, tumors were annotated for metastatic status, site of spread, grade, and Ki-67 index (**Figures 2F-2I**). As expected, metastatic cases clustered toward higher histologic grades and higher Ki-67 percentages, and most distant recurrences involved the lymph node, with a smaller subset showing liver or multi-organ dissemination. Collectively, these data indicate that while classical clinicopathologic markers (grade, Ki-67, and site of spread) stratify aggressive behavior, metastatic propensity in low-grade pNETs is not simply explained by gross differences in mutational load or base-substitution patterns, motivating deeper pathway-level analyses.

### Recurrently mutated genes and canonical cancer drivers in non-metastatic and metastatic pNETs

To delineate gene-level differences between indolent and aggressive disease, we next mapped somatic alterations across all the primary pNETs using oncoprint representations of the top 50 recurrently mutated genes together with seven canonical cancer-driver genes (**Figure 3** and **Supplementary Figure 1**). All tumors (including 56 non-metastatic and 19 metastatic) harbored at least one coding alteration in this gene set (100% altered in each cohort), underscoring the pervasive yet heterogeneous nature of the mutational landscape. Samples are ordered from left to right based on decreasing TMB, with the stacked bars above each column illustrating that missense variants comprising the majority of mutations, followed by frameshift and nonsense events. The multi-hit alterations were found to be very common in highly mutated cases.

**Figure 3.**
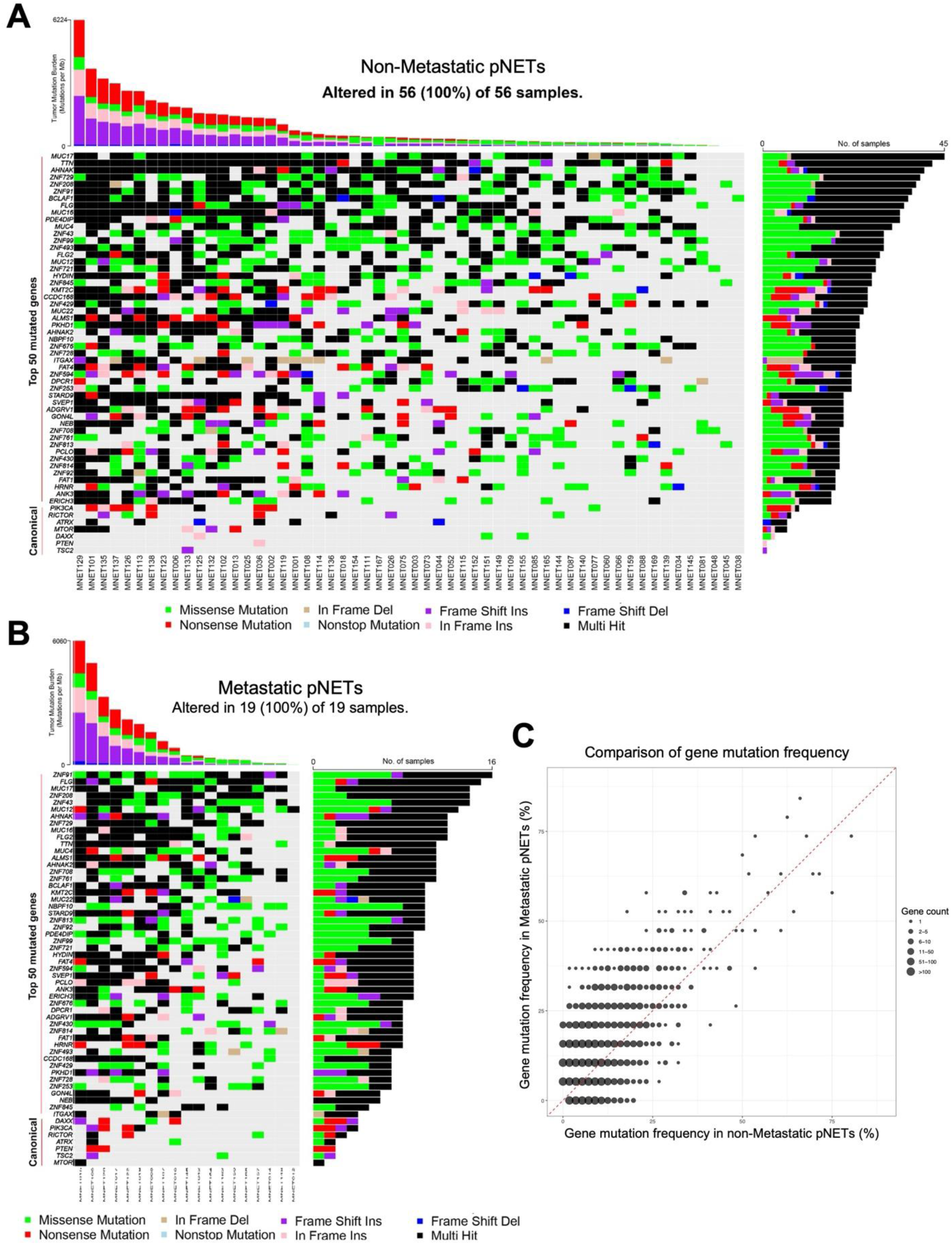
Landscape of somatic alterations and tumor mutational burden in the pNET study cohort. **A.** Oncoprint showing the top fifty recurrently mutated genes and seven canonical genes across 56 primary pNETs from the patients with non-metastatic status. **B.** Oncoprint showing the top fifty recurrently mutated genes and seven canonical genes across 19 primary pNETs from the patients with metastatic status. Each column represents an individual tumor, and each row represents a gene, ordered by decreasing mutation frequency. The stacked bar plot above the heatmap depicts tumor mutational burden (TMB; mutations per megabase) for each sample. The height of the bar reflects total TMB and the colors indicate the contribution of different mutation classes. The horizontal bar plot on the right shows the number of tumors harboring at least one mutation in each gene. Genes in the lower “Canonical” panel highlight established cancer-driver genes analyzed in this cohort. Colored boxes within the heatmap denote the mutation type observed in that gene-tumor pair: missense mutation (green), non-sense mutation (red), nonstop mutation (light pink), in-frame deletion (dark orange), in-frame insertion (beige), frameshift deletion (blue), frameshift insertion (purple), and multi-hit events (black; ≥2 mutations in the same gene in a given tumor). Grey cells indicate that no coding mutation was detected in that gene in the corresponding tumor. All variants shown are somatic single-nucleotide variants or small insertions/deletions that passed standard quality-control filters, and samples are ordered from left to right by decreasing TMB. **C.** Bubble scatterplot contrasting the prevalence of recurrent gene mutations in metastatic versus non-metastatic pNETs. The mutation frequencies are expressed as the percentage of cases harboring ≥1 coding alteration. Each bubble represents one or more genes that share the same frequency pair and bubble size encodes the number of genes at that coordinate (legend, right). The red dashed diagonal marks parity between groups.

In non-metastatic pNETs, somatic alterations were broadly distributed across a large set of genes, with frequent mutations in large structural or membrane-associated genes (e.g., MUC17, TTN, AHNAK, MUC16, FLG, MUC4, FLG2, MUC12) and numerous zinc-finger transcription factors (ZNF729, ZNF208, ZNF9, ZNF43, ZNF99, ZNF493, ZNF721 and ZNF845) among others (**Figure 3A**). Many of these genes showed multi-hit patterns in a subset of tumors, suggesting ongoing selective pressure or a mutational bias toward these loci. Canonical tumor suppressors and signaling components such as DAXX, ATRX, PIK3CA, RICTOR, PTEN, TSC2, mTOR were also recurrently altered, typically through missense, nonsense, or frameshift mutations, consistent with disruption of chromatin remodeling and PI3K-mTOR pathway signaling in a substantial fraction of non-metastatic tumors.

Metastatic pNETs displayed a qualitatively similar mutational repertoire (**Figure 3B**). The same broad classes of genes dominated the top mutated list, including mucins (MUC17, MUC16, MUC12), cytoskeletal or scaffolding genes (AHNAK, TTN), and multiple zinc-finger genes (ZNF9, ZNF43, ZNF208, ZNF729, ZNF761, ZNF813, ZNF430). As in the non-metastatic group, these loci frequently carried multi-hit events, particularly in tumors with the highest TMB. Canonical drivers in chromatin remodeling and PI3K-mTOR signaling were again recurrently mutated across metastatic samples, indicating that these pathways are commonly perturbed. Comparison of the right-hand barplots, which summarize the number of tumors mutated in each gene, revealed extensive overlap in the most frequently altered genes between non-metastatic and metastatic cohorts, with modest shifts in rank order for metastasis-specific drivers. Overall, these data show that both non-metastatic and metastatic low-grade pNETs share a complex but largely conserved set of recurrent somatic alterations, centered on mucin/structural genes, zinc-finger transcription factors.

To complement the oncoprints, we contrasted gene-level mutation prevalence between metastatic and non-metastatic pNETs using a bubble scatterplot (**Figure 3C**). Most bubbles fall near the red dashed diagonal, indicating broad parity in mutation prevalence across disease states. Nevertheless, a subset of bubbles deviates above the diagonal consistent with genes showing relatively higher mutation frequency in metastatic tumors. Such frequency-shift signals in **Figure 3C** point to a limited set of genes that may modulate metastatic propensity. The dispersion pattern, together with the oncoprints, supports a largely conserved mutational backbone with selective frequency shifts in a limited gene subset rather than wholesale rewiring of the gene-level landscape. These observations suggest that additional layers such as transcriptional programs, pathway activity are likely required to explain metastatic divergence.

### Convergent disruption of canonical oncogenic pathways and enriched proliferative programs in metastatic pNETs

We next examined how the somatic mutational landscape in pNETs converges on established oncogenic signaling pathways (**Figure 4**). To assess such biological relevance of the observed mutations, we performed pathway enrichment analysis using the Maftools mutation annotation framework. This analysis revealed significant enrichment in several oncogenic and developmental pathways, including RTK-RAS signaling, WNT/β-catenin, NOTCH, Hippo, cell cycle regulation, TGF-β signaling, PI3K/mTOR, Myc, and p53 pathways (**Figure 4A-4B**). Notably, the mutated genes in the RTK-RAS, WNT, NOTCH, and Hippo pathways were found to be activated in pancreatic cancer tissues^30^, suggesting that these alterations may contribute to pNET tumorigenesis through shared oncogenic mechanisms. Thus pathway-level aggregation revealed that virtually all tumors harbor alterations in one or more canonical cancer pathways. Although a subset of the many genes annotated to each pathway were mutated, these alterations collectively affected the pathway deregulation rather than single-gene lesions which could be a defining feature of pNET genomics. Cell-cycle regulators, MYC-related signaling, TP53, TGF-β, and NRF2 stress-response components were also recurrently affected, supporting broad involvement of growth-control, differentiation, and stress-adaptation modules in pNETs.

**Figure 4.**
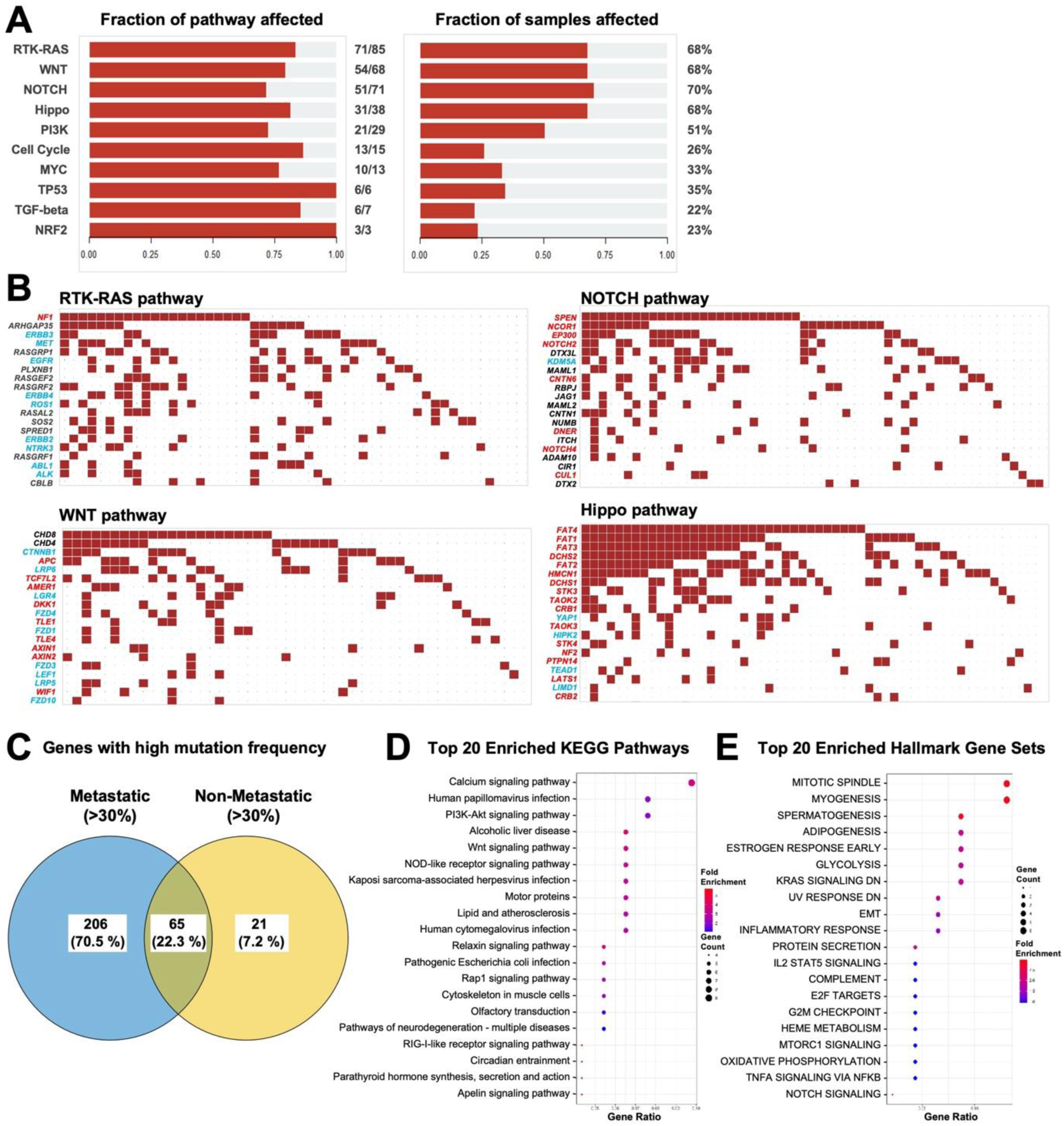
Canonical oncogenic pathway alterations and functional enrichment in hypermutated pNETs. **A.** Pathways impacted by the changes in mutational landscape in pNETs compared to normal pancreas tissues. Bar plots summarize the distribution of somatic alterations across major oncogenic signaling pathways. Pathways are ordered by the decreasing number of altered genes. For each pathway, the relative fraction of alterations among all pNET samples is shown in the right panel. **B.** Gene-level mutation maps for canonical signaling pathways including RTK-RAS, NOTCH, WNT and Hippo signaling. Columns represent individual tumors and rows represent individual genes within each pathway. The red squares indicate the presence of at least one somatic mutation in that gene in the corresponding tumor. Empty cells denote no detectable coding alteration. Gene names in red mark core canonical drivers, whereas those in blue highlight essential signaling or other potentially druggable components. **C.** Venn diagram depicting the overlapping mutated gene numbers between non-metastatic and metastatic patient pNET samples. Percentages representing the proportion of the combined gene set. **D.** Functional enrichment of top twenty KEGG pathways in metastatic patient pNET samples as compared to non-metastatic samples. **E.** Functional enrichment of top twenty hallmark gene sets in metastatic patient pNET samples as compared to non-metastatic samples. Each dot corresponds to a significantly enriched pathway or gene-ontology term. The x-axis denotes enrichment significance, the color scale indicates fold-enrichment, and dot size reflects the number of genes from the input list contributing to each term.

At the gene level, binary mutation maps for RTK-RAS, NOTCH, WNT, and Hippo signaling highlighted extensive inter-tumoral heterogeneity with clear evidence of pathway convergence (**Figure 4B**). Within each pathway, somatic mutations were distributed across multiple receptors, core signal transducers, and transcriptional effectors. Core canonical drivers (highlighted in red) and essential, often druggable, signaling nodes (highlighted in blue) were mutated in overlapping but non-identical subsets of tumors, indicating that distinct combinations of lesions can perturb the same signaling axis. Hypermutated tumors frequently carried multiple hits within a given pathway, whereas lower-burden tumors tended to harbor single-gene alterations, suggesting quantitative differences in pathway disruption while maintaining qualitative convergence on the same networks.

To relate these alterations to metastatic behavior, we compared the sets of mutated genes between non-metastatic and metastatic pNETs. Of all genes mutated in at least one tumor, 65 (22.3%) were shared between the two groups, whereas 206 (70.5%) and 21 (7.2%) were uniquely mutated in non-metastatic and metastatic tumors, respectively (**Figure 4C**). Thus, although the majority of mutated genes are restricted to metastatic tumors, a distinct subset of genes is preferentially altered in non-metastatic disease, superimposed on a common core of shared lesions. Functional enrichment of genes mutated in metastatic versus non-metastatic tumors revealed significant over-representation of multiple KEGG pathways such as Calcium, PI3K-Akt, WNT etc. signaling pathways linked to growth signaling, cell-matrix interaction, and intracellular communication (**Figure 4D**), as well as hallmark gene sets such as mitotic spindle, E2F targets, KRAS signaling etc. associated with cell-cycle progression, MYC activity, DNA damage responses, and stress-adaptation programs (**Figure 4E**). NOTCH and WNT signaling pathways are also found to be enriched in grade 2 metastatic pNETs (**Supplementary Figure 2**). Together, these data indicate that pNETs, and particularly hypermutated and metastatic tumors, display substantial gene-level heterogeneity but converge on a limited number of canonical oncogenic pathways and proliferative programs that may underlie metastatic competence and provide rational targets for therapeutic intervention.

### Transcriptomic reprogramming and metastasis-associated gene expression programs in low-grade pNETs

We first compared the transcriptomes of low-grade pNETs with normal pancreas to define tumor-specific gene expression programs (**Figure 5**). The volcano plot revealed a substantial transcriptional rewiring, with 3,187 genes significantly upregulated and 1,363 genes significantly downregulated in tumors relative to normal tissue (**Figure 5A**). Among the most strongly downregulated genes were the immune and signaling regulators MKNK1-AS1, SOS2, CXCL10, TRAT1, and TRBV17, whereas highly upregulated transcripts included SNTG1, RIMS2, GPC6, AL355255, and PTPRT, highlighting coordinated suppression of selected inflammatory/signal-transduction modules and induction of genes implicated in synaptic signaling, adhesion, and receptor regulation. The differential expression pattern was consistent when compared between non-metastatic and metastatic categories in Grade 1, Grade 2 or all pNETs (**Supplementary Figure 3**).

**Figure 5.**
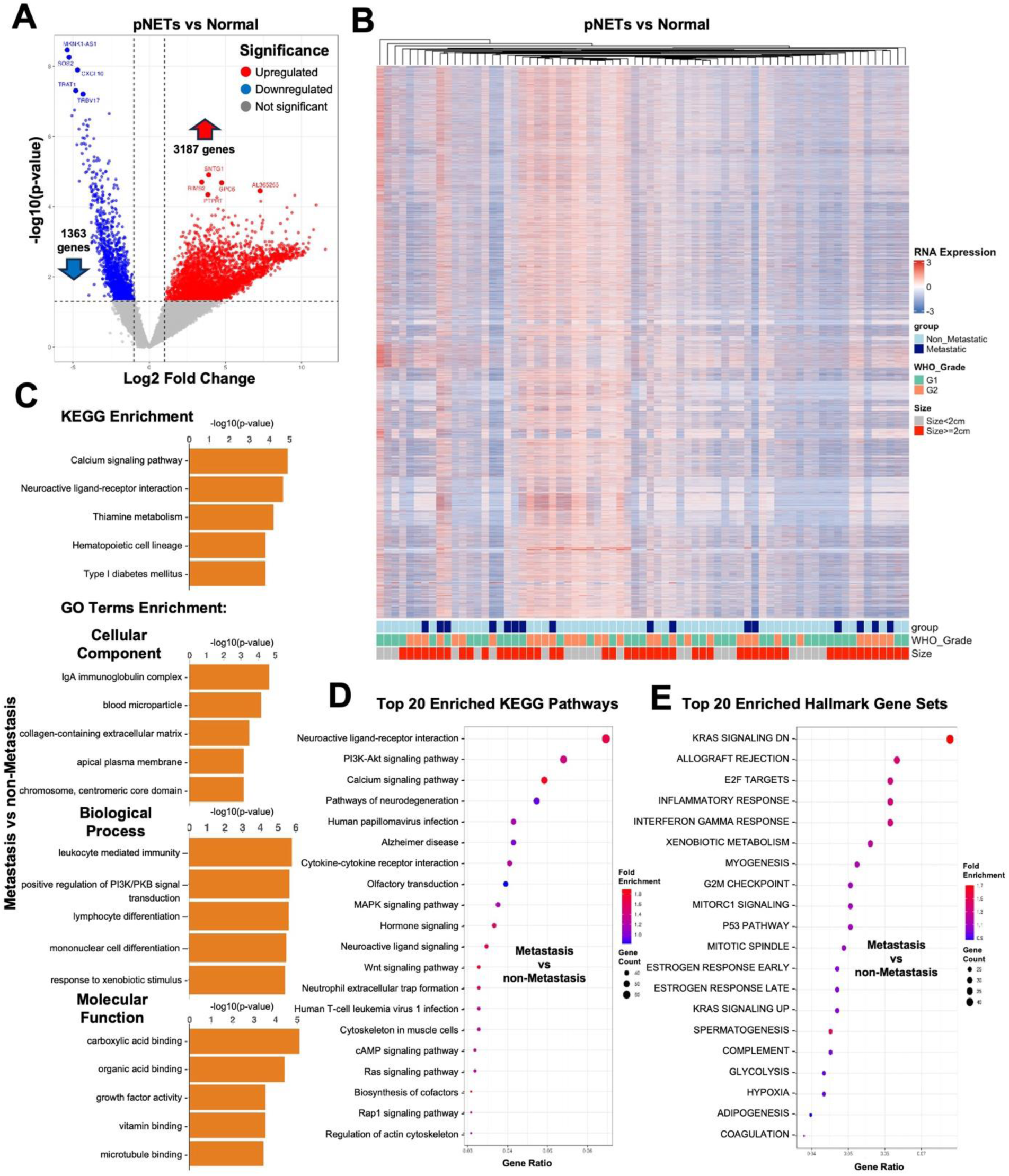
Differential gene expressions in low-grade pNET patient samples. **A.** Volcano plot of differentially expressed genes in pNET patient samples compared to normal pancreas. The x-axis shows log2 fold-changes and the y-axis shows -log₁₀ adjusted *P* value. Red and blue points indicate significantly upregulated and downregulated genes, respectively. The top -up and - downregulated genes are labelled. **B.** Unsupervised hierarchical clustering heatmap of differentially expressed genes (DEGs). Each column represents an individual tumor and each row a gene. Relative expressions depicted in a color scale where red indicates higher expression and blue indicates lower expression. The annotation bars below the heatmap indicate clinical/pathologic features for each sample: Group, metastatic status, WHO grade (G1 and G2) and tumor size (≤2 cm vs >2 cm). **C.** Horizontal bar plots showing the functional enrichment analysis of DEG between non-metastatic and metastatic tumors. The top KEGG and Gene Ontology (GO; biological process, cellular component and molecular function) enrichments are ranked by significance. **D.** DEG associated functional enrichment of top twenty KEGG pathways in metastatic patient pNET samples as compared to non-metastatic samples. **E.** DEG associated functional of top twenty hallmark gene sets in metastatic patient pNET samples as compared to non-metastatic samples. Each dot corresponds to a significantly enriched pathway or GO term. The x-axis denotes enrichment significance, the color scale indicates fold-enrichment, and dot size reflects the number of genes from the input list contributing to each term.

Unsupervised hierarchical clustering of all differentially expressed genes (DEGs) segregated tumors into distinct expression clusters (**Figure 5B**). Although clusters were not perfectly concordant with any single clinicopathologic variable, the annotation tracks demonstrated that metastatic status, WHO grade, and tumor size were not randomly distributed across the dendrogram. Metastatic and WHO Grade 2 tumors, and those >2 cm, tended to aggregate within expression subgroups. These findings suggest that discrete transcriptomic states underlie a spectrum of biological behavior within histologically low-grade pNETs.

To explore functional consequences of these transcriptional changes, we performed KEGG pathway and Gene Ontology enrichment analyses on DEGs between non-metastatic and metastatic tumors (**Figure 5C**). The top enriched KEGG pathways and GO biological process, cellular component, and molecular function terms converged on programs related to cell-cycle regulation, DNA replication and repair, extracellular matrix organization, cell-cell/cell-matrix interaction, and cytokine/immune signaling, indicating that metastatic tumors are preferentially enriched for proliferative, matrix-remodeling, and microenvironment-interacting gene sets.

Gene-set enrichment using curated KEGG and hallmark signatures further highlighted these differences (**Figure 5D-5E**). In metastatic versus non-metastatic tumors, significantly enriched pathways showed higher fold-enrichment and larger contributing gene counts for multiple proliferative, stress-response, and signaling programs, including PI3K-Akt, Calcium, MAPK, WNT, RAS signaling pathways in KEGG (**Figure 5D**) and KRAS signaling, E2F targets, mitotic spindle in hallmark gene sets are consistent with mutation associated modulations. Together, these data demonstrate that, despite their low histologic grade, pNETs exhibit extensive transcriptional reprogramming relative to normal pancreas, and that metastatic competence is associated with a distinct enrichment of gene expression programs supporting proliferation, stress adaptation, and tumor microenvironment crosstalk.

### Integrated genomic-transcriptomic analysis identifies Calcium, WNT, and KRAS/PI3K signaling as convergent axes in low-grade pNETs

To pinpoint alterations that are both genetically and transcriptionally encoded in the same tumors, we integrated somatic mutation and RNA-seq data from the low-grade pNET cohort (**Figure 6**). Intersection of significantly mutated genes with DEGs revealed a relatively small but potentially high-value overlap. A total of 176 genes (3.4%) were exclusively mutated, 4,902 genes (96.0%) were exclusively differentially expressed, and only 29 genes (0.6%) were both significantly mutated and differentially expressed (**Figure 6A**). Thus, while pNETs display broad transcriptional reprogramming, a restricted subset of genes shows concordant DNA- and RNA-level deregulation. The lower panel of **Figure 6A** shows that these 29 overlapping genes are mutated at variable frequencies across non-metastatic and metastatic tumors. Among the overlaps, zinc finger protein 273 gene ZNF273, intracellular calcium release channel gene ryanodine receptor 1 (RYR1) showed highest mutation prevalence in metastatic cases.

**Figure 6.**
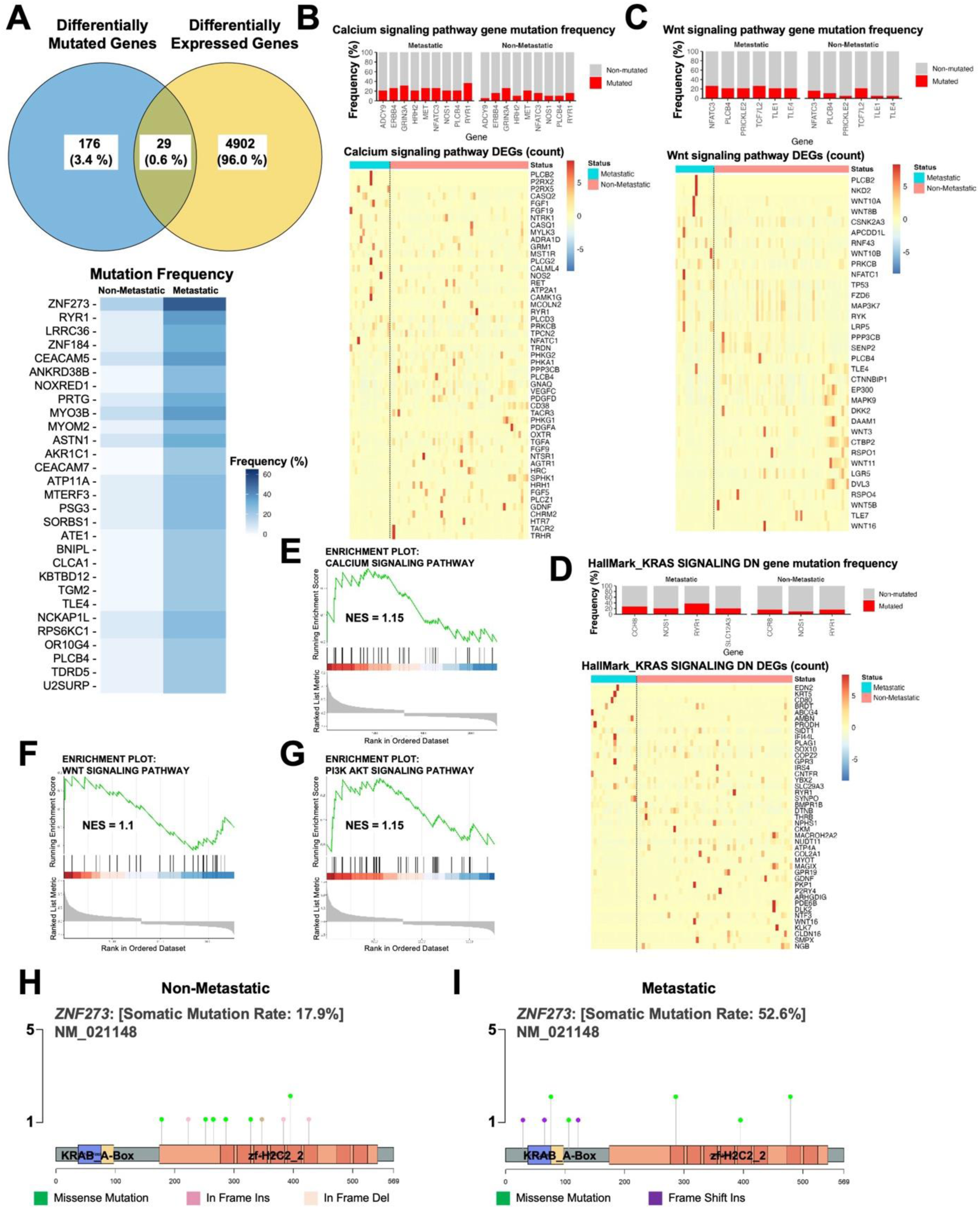
Integration of mutational alterations and transcriptional changes derived from same tumors highlights key genes and pathways in low-grade pNETs. **A.** Venn diagram showing the overlap between significantly mutated genes and differentially expressed genes (DEGs) in low-grade pNETs. Number of genes are indicated in each segment. Mutational frequency of overlapping genes in non-metastatic and metastatic pNET samples are shown in the lower panel. **B-C.** Key pathways associated with overlapped genes. Upper panel indicates gene mutation frequencies both in non-metastatic (right) and metastatic (left) pNET samples. Lower heatmap indicates the count of DEGs in non-metastatic (pink) and metastatic (blue) pNET samples. Columns represent samples and rows represent genes. **B.** Calcium signaling pathway. **C.** WNT signaling pathway. **D.** Hallmark KRAS-signaling-DN gene set and association with overlapped genes. Upper panel indicates gene mutation frequencies both in non-metastatic (right) and metastatic (left) pNET samples. Lower heatmap indicates the count of DEGs in non-metastatic (pink) and metastatic (blue) pNET samples. Columns represent samples and rows represent genes. **E-G.** Gene set enrichment analysis (GSEA) of calcium signaling, WNT signaling and PI3K-AKT signaling pathways. The enrichment plot shows normalized enrichment score (NES) for the KEGG pathways in metastatic pNETs compared to non-metastatic tumors as a function of the ranked gene list. **H.** Lollipop plot of ZNF273 (NM_021148) somatic mutations in non-metastatic primary pNETs (n = 56). The linear schematic depicts the ZNF273 protein with the N-terminal KRAB_A-Box domain and C-terminal zinc-finger region (zf-H2C2_2). Each lollipop represents a somatic variant identified by whole-exome sequencing, plotted at its codon position. **I.** Corresponding lollipop plot for metastatic primary pNETs (n = 19). Domain structure and scaling are identical to panel H.

Pathway mapping of the overlapping gene set highlighted Calcium signaling, WNT signaling, and KRAS/PI3K-related programs as major convergent nodes. In the Calcium signaling pathway, multiple genes exhibited somatic mutations in both non-metastatic and metastatic tumors, with some genes mutated more frequently in metastatic samples (**Figure 6B**, upper panel). The accompanying heatmap (**Figure 6B**, lower panel) shows that many of these same genes are also differentially expressed, with distinct expression patterns between metastatic and non-metastatic tumors, supporting functional perturbation of calcium-dependent signaling in clinically aggressive disease. Calcium signaling along with motor proteins also highlighted when non-metastatic and metastatic pNETs of Grade 2 were compared (**Supplementary Figure 4**). A similar pattern was observed for the WNT pathway, efferocytosis, inflammatory response and E2F targets (**Figure 6C** and **Supplementary Figure 5**) genes carried recurrent mutations across the cohort, and several showed consistent expression shifts between metastatic and non-metastatic tumors, suggesting that these signaling dysregulation arises through a combination of genetic lesions and transcriptional remodeling.

We next focused on the Hallmark KRAS signaling DN gene set, which encompasses downstream targets suppressed upon oncogenic KRAS activation. Several components of this gene set were mutated at appreciable frequencies in both metastatic and non-metastatic tumors (**Figure 6D**, upper panel), and the corresponding heatmap (**Figure 6D**, lower panel) demonstrated coherent expression changes across the cohort. These data imply that KRAS-related signaling, although not always driven by classic KRAS mutations, is modulated through alterations in its downstream effector network.

To assess whether these pathways are coordinately enriched at the transcriptomic level in metastatic disease, we performed gene set enrichment analysis (GSEA) comparing metastatic versus non-metastatic tumors. Calcium signaling, WNT signaling, and PI3K-AKT signaling were all positively enriched in metastatic pNETs (**Figure 6E-6G**), with normalized enrichment scores (NES) above 1, indicating a consistent, cohort-wide shift toward higher pathway activity in metastatic pNETs. Collectively, these integrative analyses demonstrate that only a small fraction of genes are both mutated and differentially expressed, but those that tend to cluster within a limited set of signaling pathways most notably Calcium, WNT, and KRAS/PI3K-AKT that are transcriptionally upregulated in metastatic tumors. These convergent axes likely represent key molecular circuits underpinning metastatic competence in low-grade pNETs and may provide rational targets for therapeutic intervention or biomarker development.

We have analyzed mutational patterns for top overlapping genes including ZNF273, CLCA1, RYR1, CEACAM5, MYO3B and observed distinct distribution of mutations both in percentage and types between non-metastatic and metastatic pNET samples (**Figure 6H-6I** and **Supplementary Figure 6**). Lollipop plots of ZNF273 (NM_021148) in pNETs reveal a marked increase in somatic alterations in metastatic versus non-metastatic tumors. In non-metastatic cases (n = 56), ZNF273 is mutated in 17.9% of tumors, primarily through missense substitutions and a smaller number of in-frame insertions and deletions distributed across the C-terminal zinc-finger region (zf-H2C2_2) and, less frequently, the N-terminal KRAB_A-box domain. In contrast, metastatic primaries (n = 19) show a substantially higher somatic mutation rate of 52.6%, with multiple recurrent missense variants and additional frame-shift insertions affecting both the KRAB_A-box and zinc-finger domains, indicating more disruptive lesions in the transcriptional regulatory architecture of ZNF273. Collectively, these data demonstrate that metastatic pNETs are enriched for structurally diverse and potentially more deleterious ZNF273 mutations compared with non-metastatic tumors, consistent with a role for ZNF273 dysregulation in metastatic competence.

### In-silico drug prioritization reveals candidate targeted and repurposable therapies for metastatic low-grade pNETs

To explore whether the molecular alterations identified in metastatic low-grade pNETs might be pharmacologically tractable, we applied complementary pathway- and gene-centric drug prediction strategies (**Figure 7**). Upstream regulator analysis using iPathwayGuide, which integrates the direction and magnitude of DEGs with a curated drug-target interaction network, nominated several compounds whose known targets were consistently perturbed in metastatic versus non-metastatic tumors (**Figure 7A**). These agents showed large numbers of “consistent DEGs” (bars to the right), indicating that the observed transcriptional changes in metastatic pNETs would be expected to be reversed or attenuated by drug exposure, and therefore may represent clinically actionable candidates in this setting. In contrast, a second group of drugs was predicted to lack actionable changes or to drive gene-expression shifts discordant with those observed in metastatic tumors based on similarly structured analyses (**Figure 7B**). These compounds, which also showed substantial DEG coverage, may be less likely to provide benefit in metastatic low-grade pNETs and could even be counter-therapeutic.

**Figure 7.**
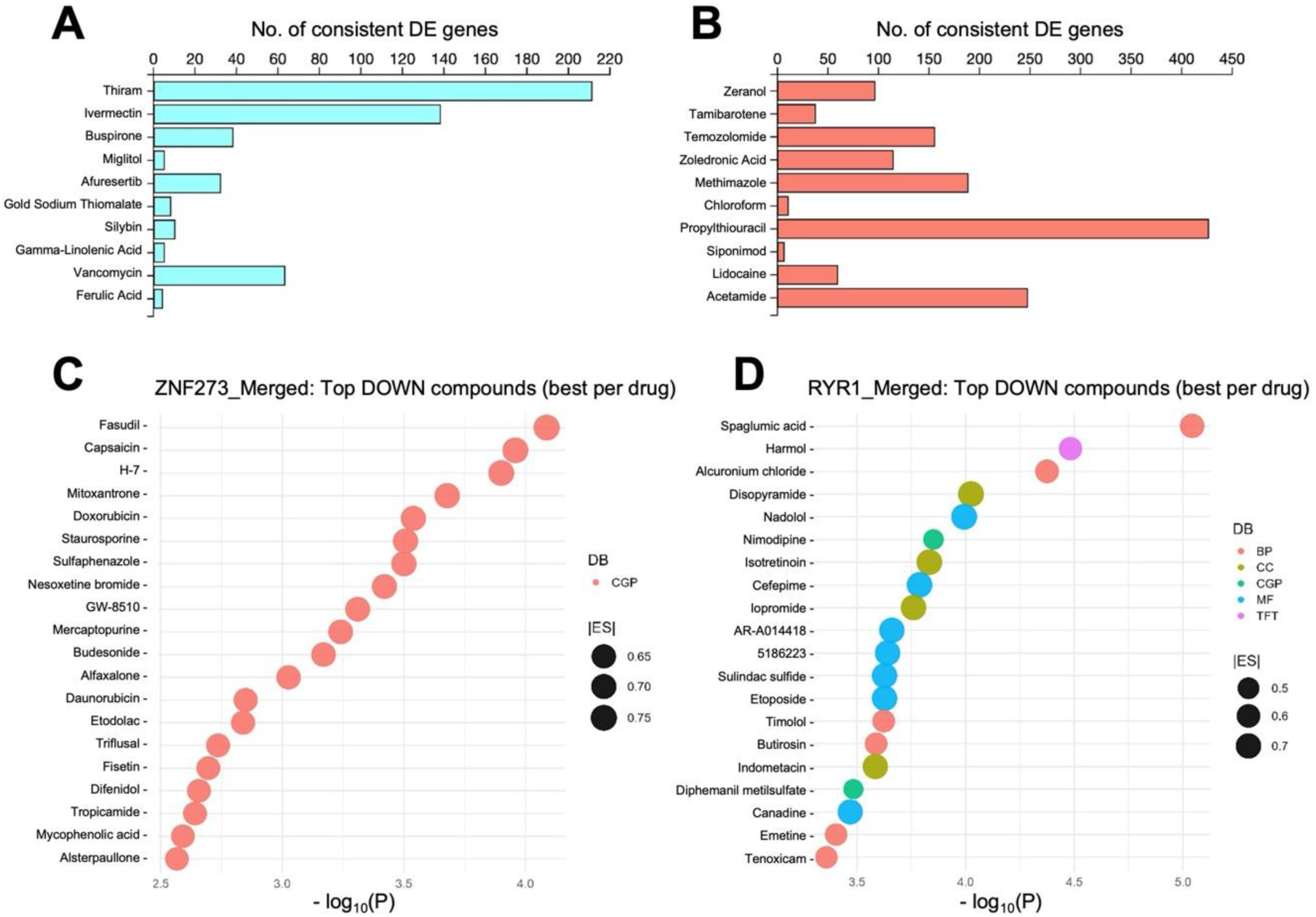
Clinically actionable genetic alterations observed in low-grade pNETs with metastatic status. **A.** iPathwayGuide analysis (Advaita Corp. 2025) of gene expressions in metastatic pNETs compared to non-metastatic pNETs predicted drugs that could have actionable changes in low-grade pNETs that have metastasized in the patients. The prediction of upstream drugs is based on the enrichment of differentially expressed genes (DEGs) from the experiment and a network of interactions from the Advaita Knowledge Base (AKB v18.1). Bar diagram indicates number of consistent DEGs. **B.** iPathwayGuide analysis of gene expressions in metastatic pNETs compared to non-metastatic pNETs predicted drugs that may not have actionable changes in low-grade pNETs have metastasized in the patients. Bar diagram indicates number of consistent DEGs. **C-D.** The Gene2Drug software predicted multiple drugs against topmost overlapped genes (after integration of genetic alteration and gene expression data) observed in the study having high tumor mutational burden in metastatic pNETs. DOWN indicates inhibitory effect. **C.** Gene2Drug-predicted drugs against ZNF273 alterations. **D.** Gene2Drug-predicted drugs against RYR1 alterations. BP, biological pathway; CC, cellular component; CGP, cancer-related drug perturbation data; DB, database; ES, enrichment score; MF, molecular function; TFT, Transcription-factor-target sets.

We next used Gene2Drug software to focus on specific genes that were both recurrently altered and associated with high TMB in metastatic tumors following integration of exome and RNA-seq data. This analysis highlighted ZNF273 and RYR1 as representative “overlapped” genes whose perturbation may be central to the metastatic phenotype. Gene2Drug queried multiple functional databases, including cancer drug perturbation signatures (CGP) and Gene Ontology categories (BP, CC, MF) as well as transcription-factor-target (TFT) sets, to identify drugs that reproducibly downregulate expression programs centered on these genes. For ZNF273, numerous compounds demonstrated strong inhibitory (DOWN) signatures with relatively high enrichment scores (ES; 0.65-0.75), suggesting that their transcriptional footprints oppose the ZNF273-associated expression pattern in metastatic tumors (**Figure 7C** and **Supplementary Figure 7**). A similar analysis for RYR1 yielded a partially overlapping but distinct set of candidate drugs (ES; ∼0.7) drawn from multiple annotation databases (**Figure 7D** and **Supplementary Figure 8**).

Although exploratory and requiring functional validation, these integrative analyses demonstrate that metastatic low-grade pNETs harbor molecular alterations that map onto existing pharmacologic space. By combining pathway-level upstream regulator analysis with gene-centric perturbation signatures, our data nominates a focused set of candidates targeted and repurposable agents that may be capable of modulating key transcriptional programs driven by ZNF273, RYR1, and related networks in metastatic low-grade pNETs.

## Discussion

Pancreatic neuroendocrine tumors lacking MEN1 mutations comprise a major and under characterized subset. By profiling low-grade primary pNETs with whole-exome and RNA sequencing, including 25 percent with documented metastasis, we close a key gap by showing that metastatic behavior emerges from pathway-level convergence rather than general changes in mutational burden. To our knowledge, this is the first comprehensive study integrating genomic and transcriptomic landscapes of low-grade pNETs to distinguish metastatic from non-metastatic disease. The central advance is that metastatic propensity in these histologically low-grade tumors is not explained by bulk mutational load or shifts in substitution spectra but is constrained by coordinated activation of Calcium, WNT, and KRAS/PI3K pathway programs. These are detectable at the transcriptome level and overlap with a small set of genes that are both mutated and differentially expressed, including ZNF273 and RYR1. Importantly, upstream regulator and gene-centric perturbation analyses nominate tractable candidates predicted to reverse metastatic expression states while flagging agents unlikely to benefit this population, providing a prioritized therapeutic hypothesis for prospective validation. Our data suggest that convergent, domain-specific mutational patterns in frequently mutated genes may constitute a molecular signature capable of stratifying metastatic risk in low-grade pNETs. These findings refine a MEN1 centric model of pNET tumorigenesis by shifting emphasis toward transcriptomic reinforcement of shared oncogenic pathways rather than acquisition of unique mutational processes.

While Chan et al. (2018) and other research groups characterized molecular subtypes defined by MEN1, DAXX, and ATRX mutations, their analyses primarily focused on MEN1-mutant tumors, leaving the MEN1-wild type subset (which accounts for ∼60% of pNETs) less explored^17,18,23^. Our data position MEN1-wild type low-grade pNETs as tumors in which metastatic potential arises from quantitative reinforcement of shared oncogenic circuits rather than acquisition of new mutational processes. Clinicopathologic enrichment of adverse features in Grade 2 tumors (**Figure 1D**) and a modest rise in mutation burden in metastatic cases (**Figure 2A**, **2C**) occur without broad changes in base-substitution spectra (**Figure 2E**), indicating conserved mutational mechanisms across disease states. Oncoprint analysis shows substantial overlap in recurrently altered genes between non-metastatic and metastatic tumors, particularly involving chromatin remodelers and PI3K-mTOR components (**Figure 3A-B**), while frequency skews for a limited subset of genes are evident in metastasis (**Figure 3C**). This pattern reinforces the concept that pathway activation, rather than mutational quantity, drives metastatic progression. These findings refine earlier genomic observations showing that sporadic pNETs generally harbor low to moderate TMB and recurrent mutations in chromatin-remodeling and PI3K-mTOR pathways^31,32^. Moreover, recent reviews emphasize that PI3K/AKT/mTOR signaling alone does not fully predict metastatic behavior, suggesting the involvement of broader network reinforcement^33^.

Virtually all tumors in our MEN1-wild-type, low-grade pNET cohort, carry alterations in one or more hallmark pathways with convergence on RTK-RAS, WNT, NOTCH, Hippo, PI3K-mTOR, MYC, TP53, TGF-β and NRF2 modules (**Figure 4A-4B**), while only a small fraction of genes are uniquely mutated in metastases yet those genes collectively enrich growth and communication pathways, including PI3K-Akt, WNT and Calcium signaling (**Figure 4C-4D**). Differential expressions relative to normal pancreas show broad rewiring with 3,187 upregulated and 1,363 downregulated genes (**Figure 5A**), and unsupervised clustering links specific transcriptomic states to metastatic status, grade and size (**Figure 5B**). Metastases preferentially activate E2F targets, mitotic spindles and KRAS-linked signatures together with matrix-interaction programs (**Figure 5C-5E**). These data align with studies establishing convergent pathway biology in pNETs in which diverse lesions perturb shared axes such as MEN1/DAXX/ATRX and PI3K-mTOR, and extend them by showing that, in MEN1-wild-type disease, metastatic behavior tracks with pathway intensity rather than the appearance of new categories of mutation^7,9^. The transcriptional enrichment of PI3K-mTOR and KRAS-proximal programs in metastases is concordant with the clinical activity of pathway-directed agents such as everolimus and sunitinib in advanced pNETs, supporting a rationale for biomarker-guided intensification or combination strategies in this subgroup^24,34^.

Integrating mutations and RNA profiles from the same MEN1-wild-type low-grade pNETs, we find that only a narrow fraction of genes shows concordant DNA/RNA deregulation (29 of 5,107; 0.6%), yet these concentrate within three reproducible signaling axes including Calcium, WNT, and KRAS/PI3K that are preferentially engaged in metastatic tumors (**Figure 6A-6G**). In Calcium signaling, metastatic cases carry higher mutation frequencies and coherent expression shifts across ryanodine-receptor and channel components (e.g., RYR1) with positive pathway enrichment by GSEA (**Figure 6B, 6E**), nominating calcium-dependent excitability as a metastasis-linked property in neuroendocrine epithelium. This aligns with broader evidence that Ca2+ circuit remodeling supports invasion and survival across cancers while extending such observations to pNETs^35–37^. In parallel, WNT nodes show recurrent lesions with consistent transcriptional upregulation in metastasis (**Figure 6C, 6F**), resonating with reports that WNT/β-catenin contributes to neuroendocrine tumor growth and invasiveness and is variably active in pNENs^38,39^. Finally, although canonical KRAS mutations are uncommon in pNETs relative to pancreatic ductal adenocarcinoma (PDAC), our data indicate modulation of the Hallmark KRAS-signaling-DN program together with PI3K-AKT enrichment (**Figure 6D, 6G**), consistent with the long-standing view from genomics that pNETs frequently perturb MEN1/DAXX/ATRX and mTOR-PI3K pathways rather than RAS itself^7^. Clinically, this convergence provides a mechanistic rationale for the efficacy of pathway-directed agents everolimus and sunitinib improved progression-free survival in randomized trials and suggests combination strategies that co-target PI3K/mTOR with WNT or Ca2+ effectors could be prospectively prioritized in metastatic, MEN1-wild type disease^14,15^. Together, these findings argue that metastatic competence in low-grade pNETs reflects quantitative reinforcement of a shared circuitry rather than wholesale pathway rewiring, and they identify calcium, WNT, and KRAS/PI3K signaling as tractable hubs for biomarker development and therapeutic testing (**Figure 6A-6G**).

Our findings identify ZNF273 and ZNF184 as a previously unappreciated Krüppel-associated box (KRAB)-zinc finger axis linked to metastatic competence and poor outcome in pNETs, extending prior genomic studies that emphasized MEN1, DAXX, ATRX and mTOR pathway alterations but did not highlight KRAB-ZNFs as recurrent lesions^7,9,40,41^. Lollipop plots show that ZNF273 is mutated in more than half of metastatic versus fewer than one fifth of non-metastatic pNETs with metastatic cases exhibiting more disruptive missense and frameshift variants across both the KRAB_A box and C2H2 zinc-finger domains (**Figure 6H-6I**). These patterns align with pan-cancer observations that mutations targeting structurally critical zinc-finger residues can rewire transcriptional programs that promote progression^42^ and with growing evidence that KRAB-ZNFs function as context-dependent oncogenic regulators^43,44^. Our results indicate that KRAB-ZNFs may stratify metastatic risk within the otherwise genetically quiet pNET landscape and warrant functional investigation as contributors rather than incidental passengers in metastatic progression.

In this study, we used complementary pathway-level and gene-centric perturbation analyses to nominate actionable and repurposable drugs and, equally important, to flag agents predicted to be non-beneficial in metastatic disease (**Figure 7A-7D**). These results extend prior genomic characterizations that emphasized MEN1/DAXX/ATRX and PI3K-mTOR lesions in pNETs by mapping metastasis-linked transcriptional programs onto pharmacologic space rather than solely onto mutation catalogs^7,9,18^. Upstream-regulator inference with iPathwayGuide aligned large fractions of metastatic DEGs with drug targets, prioritizing compounds whose predicted effects would counter the observed expression shifts (**Figure 7A**) while identifying others that were discordant and therefore less likely to help (**Figure 7B**). This approach is methodologically supported by prior descriptions of iPathwayGuide for mechanistic pathway and regulator discovery^25^. Orthogonally, Gene2Drug queries centered on overlapped, metastasis-associated genes highlighted ZNF273 and RYR1, each linked to reproducible DOWN signatures across multiple knowledge bases with moderate to strong enrichment (ES, 0.65-0.75; **Figure 7C-7D**), consistent with Gene2Drug’s validated strategy of ranking drugs by their ability to reverse target-pathway transcriptional activity^26^. While ZNF273 is novel in pNETs, RYR1 dysregulation has been implicated in tumor progression and drug repositioning in other solid tumors and has recurrent alterations reported in neuroendocrine contexts, supporting plausibility for functional relevance in pNET biology^27,28^. Notably, our drug-prediction framework complements emerging clinical sequencing data showing pathway perturbations in metastatic low-grade pNETs and provides testable hypotheses to rationalize sensitivity and resistance, including the a priori identification of agents unlikely to benefit this subset^45^. Collectively, these data argue that metastasis in MEN1-wild type low-grade pNETs is accompanied by coherent, drug-modifiable transcriptional circuits with attention to the mixed evidence base surrounding pNET molecular subtypes and outcomes^9,18^.

By deeply characterizing MEN1-wild-type, low-grade pNETs, we show that metastasis is a property of pathway state rather than simple mutation burden. Metastatic tumors exhibit slightly higher mutation frequencies but stronger activation of Calcium, WNT, and KRAS/PI3K signaling, converging on a 29-gene set that links genomic alterations to transcriptomic remodeling. Convergent activation of such programs distinguishes metastatic from non- metastatic tumors and connects directly to rational therapeutic choices. This framework shifts emphasis from single-gene events toward network activity and provides a blueprint for biomarker-guided combination trials in a clinically important subset of pNETs.

## Acknowledgements

The authors acknowledge the contribution of Dr. Julie Boerner at the Karmanos Cancer Institute Biobanking and Histopathology Core for providing the tissue samples. The authors also acknowledge Brian Burns at the Tissue Procurement Lab Coordinator Winship Cancer Institute Cancer Tissue and Pathology Shared Resource. The authors acknowledge 2P30CA022453-39 Karmanos Cancer Institute, Cancer Center Support Grant. Funding from SKY Foundation Inc., Partners Funds and U Can-Cer Vive to Dr. Azmi is acknowledged.

## Author contributions

ASA, BFE designed the study, wrote and revised the manuscript. MHU, ZM, IM, BRH, analyzed the data, wrote and revised the manuscript. YL, YS, YW, VO, GD, guided the statistical analyses. BR, HYK, AA, SFB, HJ, AJ, MNA, IA, AM, TH, NV, RB, MT, EWB, HC, AFS, PAP, JB, RMM, BCP edited and revised the manuscript.

## Data availability statement

The genetic variant data from WES of the pNET tissue samples are available from EBI’s (European Bioinformatics Institute) BioStudies repository under accession number S-BSST1766 (URL: https://www.ebi.ac.uk/biostudies/studies/S-BSST1766?key=0a84424e-187f-4c36-bc01-2d59ca9669b1). The RNA-Seq data from the FFPE and fresh pNET and adjacent normal tissues are available from EBI’s ArrayExpress repository under accession number E-MTAB-14709 (URL: https://www.ebi.ac.uk/biostudies/arrayexpress/studies/E-MTAB-14709?key=3a110a96-fca4-438e-822f-5afe8aab97ca). All data generated or analyzed during this study will be made available along with code information on reasonable request for academic use.

## Ethics approval and consent to participate

The study was approved by the Institutional Review Board (IRB) of Emory University (Atlanta, GA) with the IRB number of STUDY00001739 and of Karmanos Cancer Institute (Detroit, MI) with the IRB number of 034916MP2X. Each patient provided informed consent before participating in the study. The study protocol was approved by the Ethics Committee of Emory University (FWA00005792) and Karmanos Cancer Institute Wayne State University (FWA00002460).

