## Supplemental Information for "Genomic and Transcriptomic Landscapes of MEN1-Wild-Type Low-Grade Metastatic Pancreatic NETs Uncover Key Oncogenic Drivers and Targetable Pathways" (from another version of this preprint: version 1, 2025-12-10)

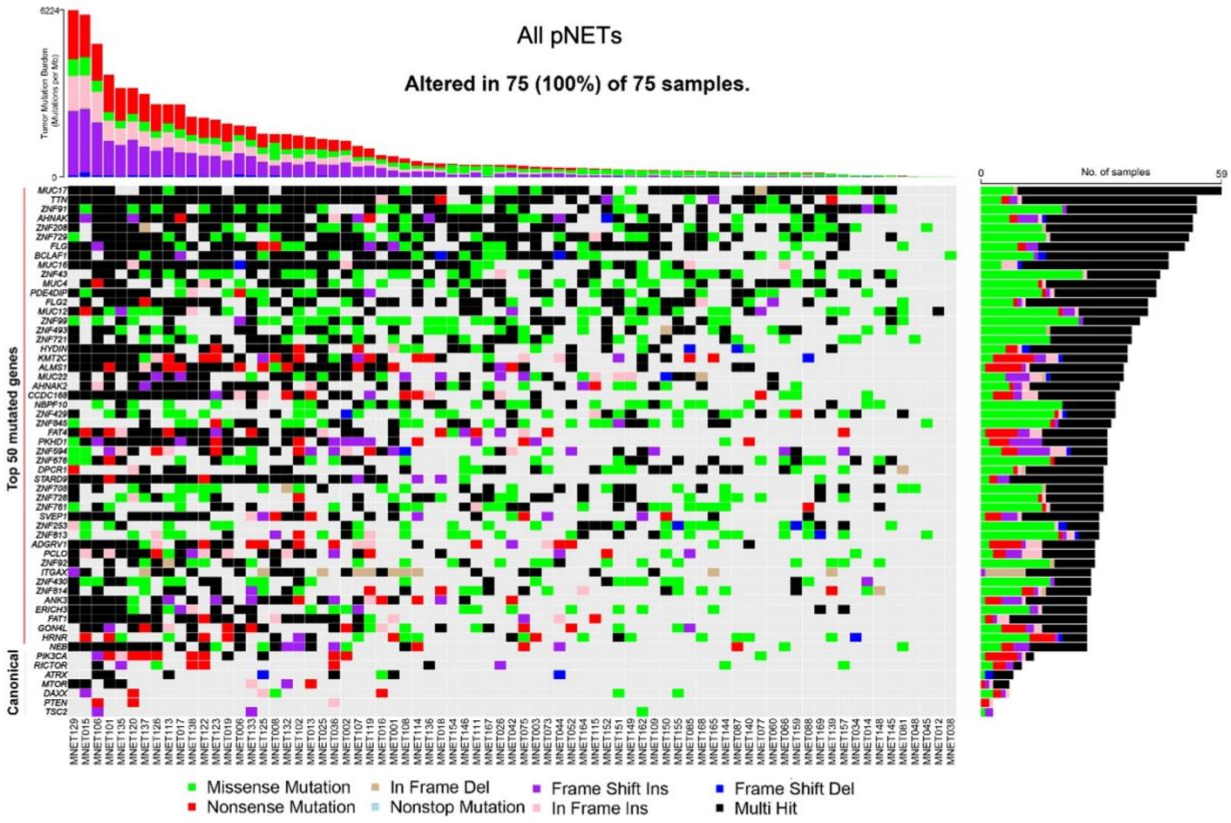


**Supplementary Figure 1. Global landscape of recurrent somatic alterations in all low-grade pNETs.** Oncoprint summarizing the top 50 recurrently mutated genes together with canonical cancer genes across all primary pNETs profiled by whole-exome sequencing (n = 75). Each column represents an individual tumor and each row a gene, ordered from top to bottom by decreasing mutation frequency. The stacked bar plot above the oncoprint depicts the tumor mutational burden (TMB) for each sample; bar height reflects overall TMB and colors indicate the contribution of different mutation classes. The horizontal bar plot to the right shows the number of tumors harboring at least one mutation in each gene. Colored boxes within the heatmap denote the mutation type identified in that gene-tumor pair: missense mutation (green), nonsense mutation (red), in-frame deletion (brown), nonstop mutation (light blue), frameshift insertion (purple), in-frame insertion (pink), frameshift deletion (blue), and multi-hit events (black). Grey cells indicate no detectable coding mutation in that gene in the corresponding tumor.

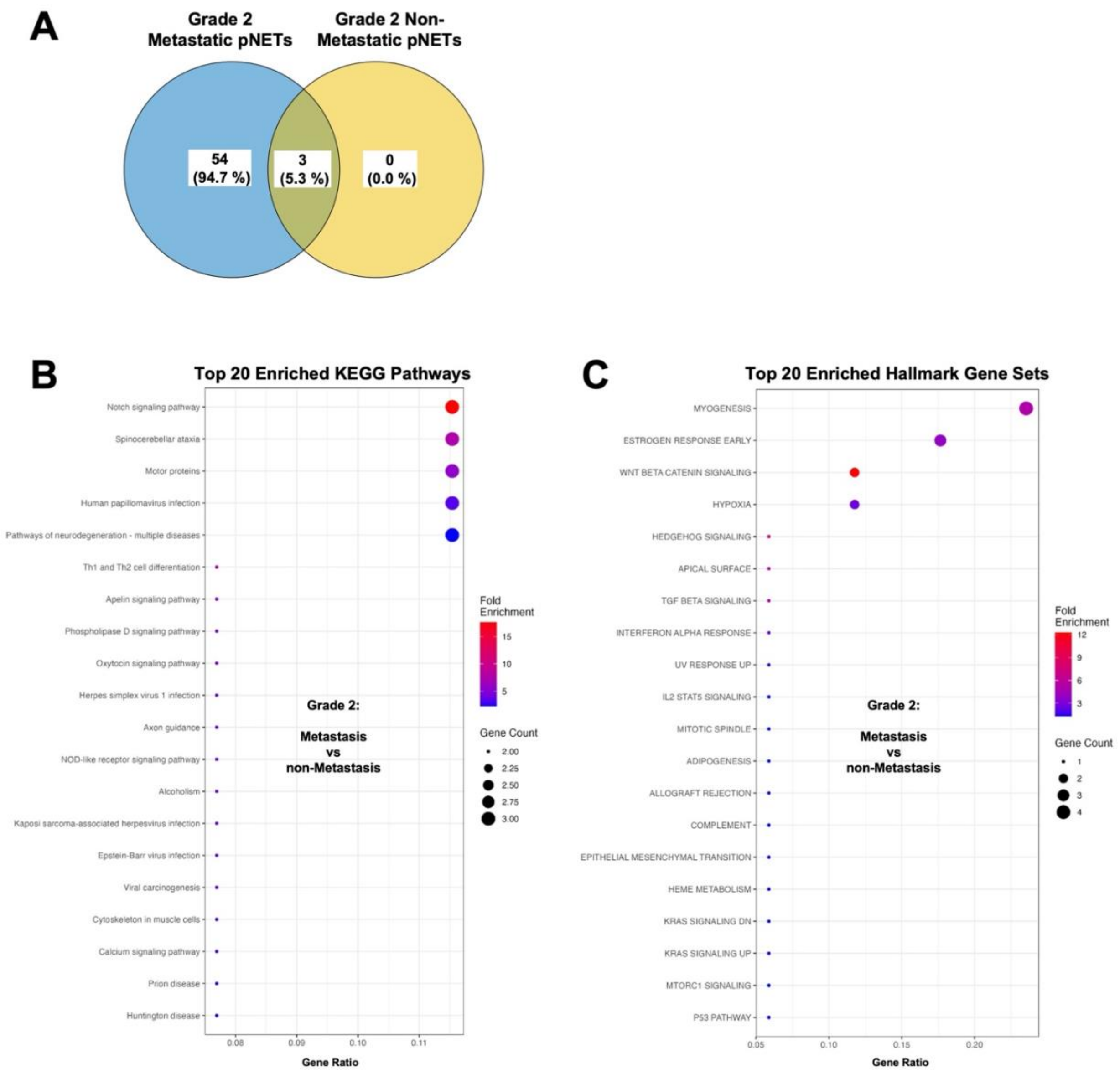


**Supplementary Figure 2. Comparison of mutational landscapes in Grade 2 metastatic versus non-metastatic pNETs. A.** Venn diagram summarizing the distribution of recurrently mutated genes in WHO Grade 2 metastatic and non-metastatic pNETs. Numbers and percentages indicate the count and proportion of genes mutated exclusively in metastatic tumors, shared between both groups, or unique to non-metastatic tumors. **B.** Dot plot showing the top 20 KEGG pathways enriched among genes mutated in Grade 2 metastatic versus non-metastatic pNETs. The x-axis depicts the gene ratio (number of mutated genes in the pathway divided by the total number of mutated genes used for enrichment). Dot size encodes the number of genes contributing to each pathway, and color represents fold enrichment, with warmer colors indicating stronger enrichment.

**C.** Dot plot of the top 20 enriched Hallmark gene sets derived from the same comparison (Grade 2 metastatic versus non-metastatic pNETs). Axes and symbol encodings are as in panel B.

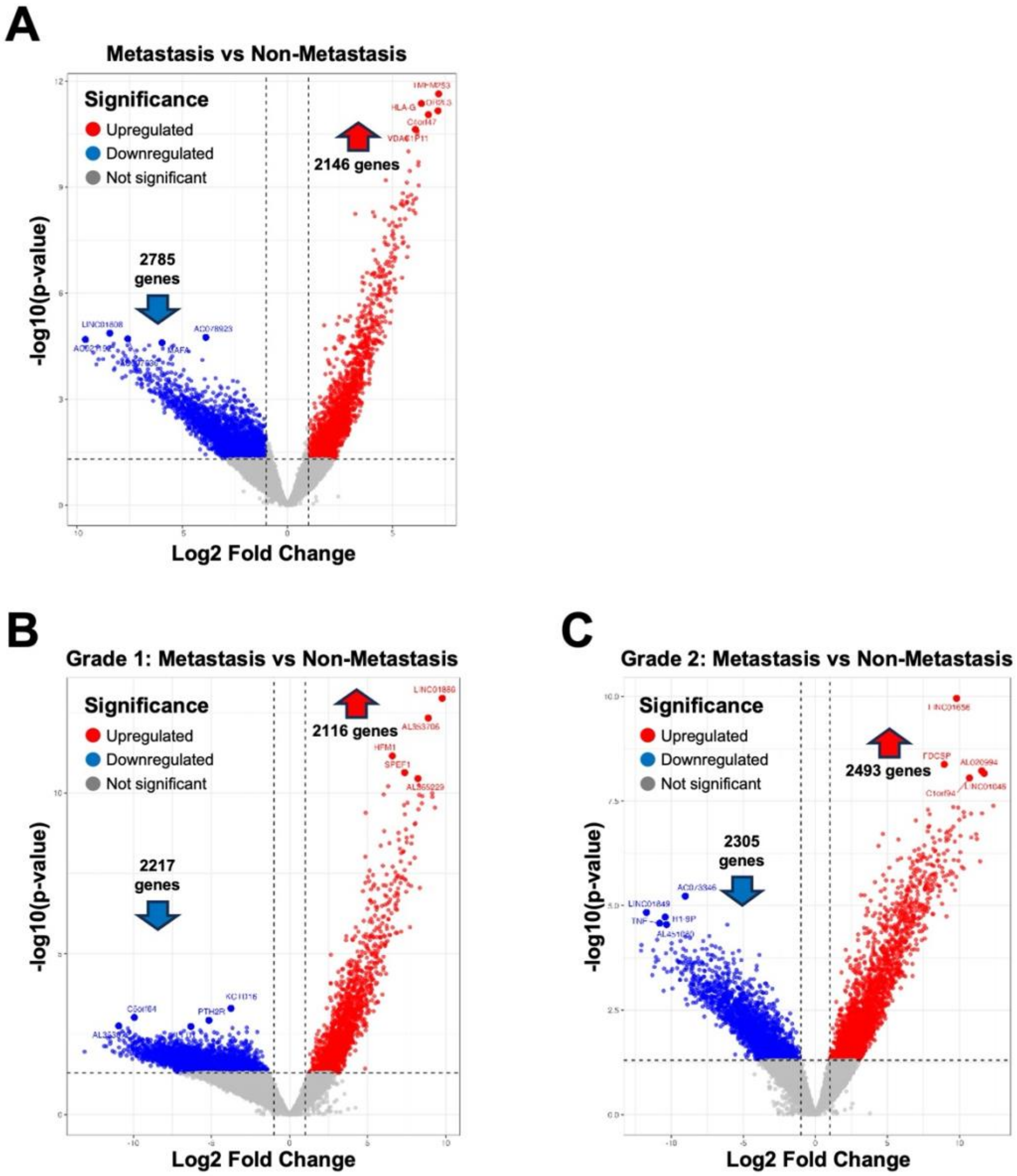


**Supplementary Figure 3. Differential gene-expression profiles associated with metastatic status in low-grade pNETs. A.** Volcano plot depicting differentially expressed genes (DEGs) between all metastatic and all non-metastatic pNETs. The x-axis shows $\log_2$(fold change) for metastatic versus non-metastatic tumors and the y-axis shows $-\log_{10}$(*p*-value). Significantly upregulated genes in metastatic tumors are shown in red, significantly downregulated genes in blue, and non-significant genes in grey (significance threshold defined by the vertical and horizontal dashed lines). Arrows indicate the total number of significantly upregulated (2,146 genes, red arrow) and downregulated (2,785 genes, blue arrow) transcripts, with representative top-ranked genes labeled. **B.** Volcano plot of DEGs comparing Grade 1 metastatic versus non-metastatic pNETs, displayed as in panel A. A total of 2,116 genes are significantly upregulated and 2,217 genes are significantly downregulated. **C.** Volcano plot of DEGs comparing Grade 2 metastatic versus non-metastatic pNETs. Using the same axes and color coding, 2,493 genes are significantly upregulated, and 2,305 genes are significantly downregulated in metastatic tumors.

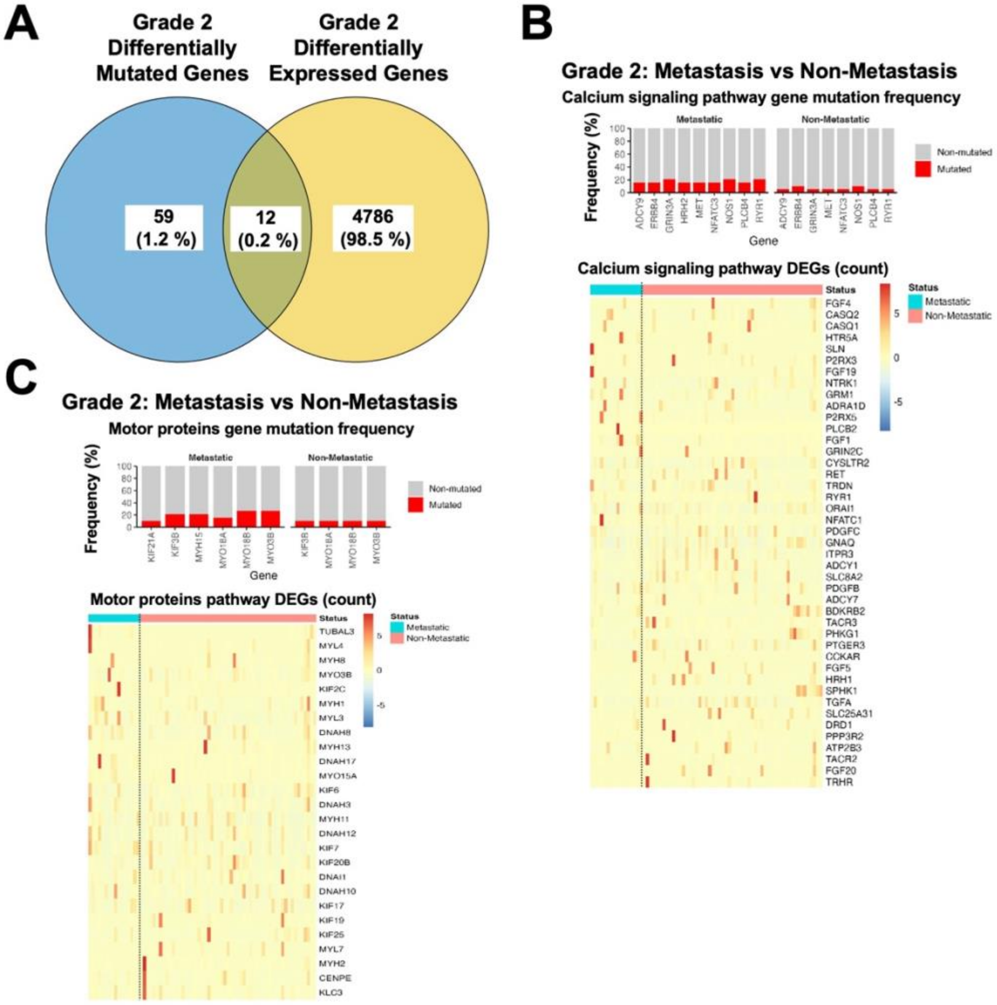


**Supplementary Figure 4. Integrated mutational-transcriptional analysis highlights calcium signaling and motor protein pathways in Grade 2 pNETs. A.** Venn diagram integrating somatic mutation and gene-expression data for all WHO Grade 2 pNETs. The blue circle denotes genes that are significantly differentially mutated between Grade 2 metastatic and non-metastatic tumors, and the yellow circle represents genes that are significantly differentially expressed (DEGs) in the same comparison. Numbers and percentages indicate the counts and proportions of genes unique to the mutational set, unique to the DEG set, or overlapping between both. **B.** Calcium signaling pathway genes identified from the overlapping set-in panel A. The upper bar plot shows the frequency of somatic mutations in each calcium pathway gene for Grade 2 metastatic and non-metastatic tumors. The lower heatmap displays the corresponding expression patterns for calcium pathway DEGs across individual tumors, with columns representing samples and rows representing genes. Expression changes are color coded with red indicating higher and blue indicating lower expression relative to the cohort mean. **C.** Motor protein pathway genes derived from the same integrated analysis. The upper bar plot depicts mutation frequencies for motor protein encoding genes in Grade 2 metastatic and non-metastatic tumors, as in panel B. The lower heatmap shows the expression profiles of motor protein pathway DEGs in the same samples, using the same color scale.

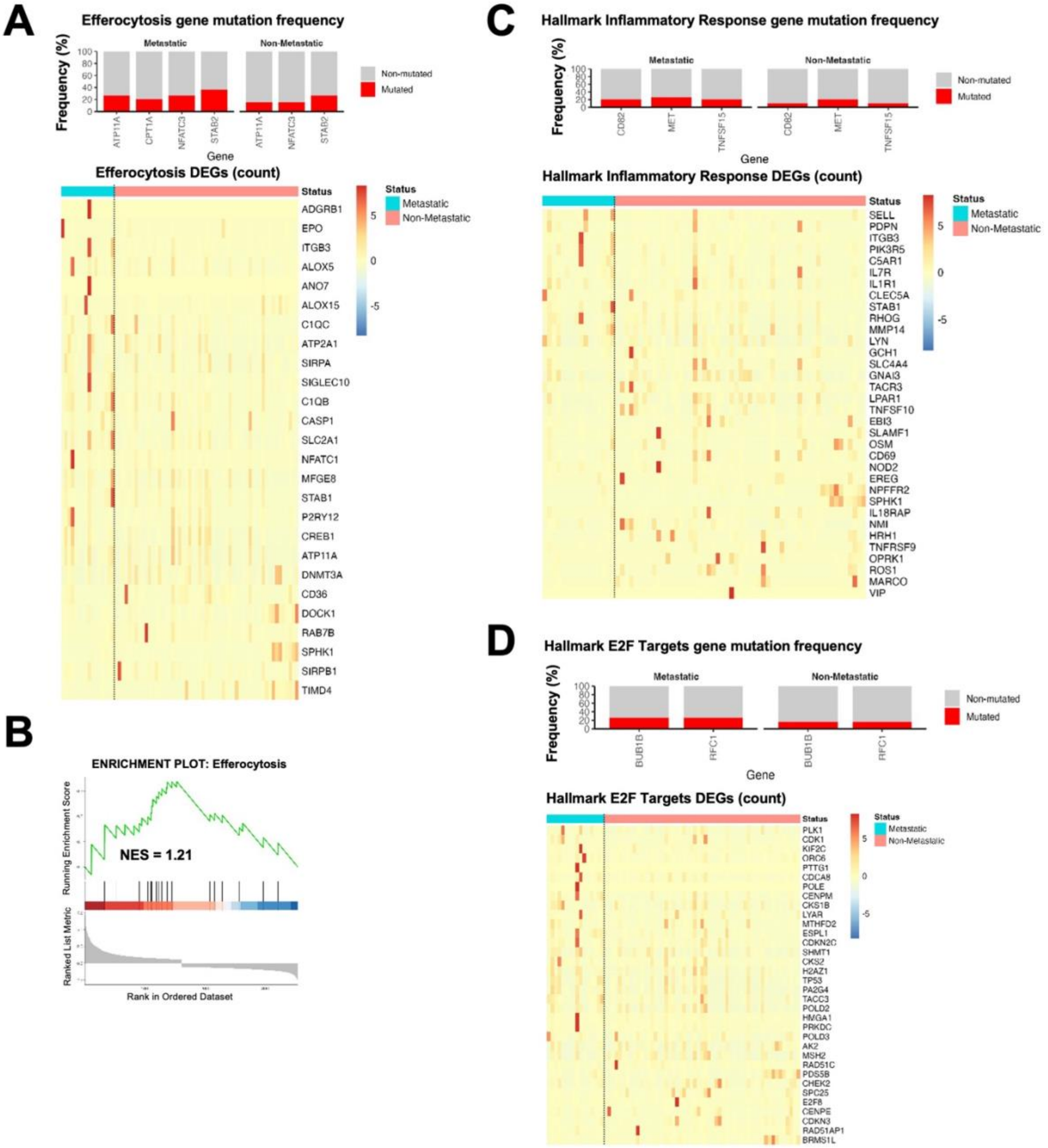


**Supplementary Figure 5. Integrative mutational and transcriptional analysis highlights efferocytosis, inflammatory response, and E2F targets programs in metastatic pNETs. A.** Efferocytosis pathway genes identified from the overlap of differentially mutated and differentially expressed genes between metastatic and non-metastatic pNETs. The upper bar plot shows, for each gene, the proportion of metastatic (left) and non-metastatic (right) tumors carrying at least one coding mutation. The lower heatmap displays expression changes for efferocytosis-related DEGs across individual tumors, with columns representing samples and rows representing genes.

Expression values are color coded, where red indicates higher and blue lower expression relative to the cohort mean. **B.** Gene set enrichment analysis (GSEA) plot for the efferocytosis pathway in metastatic versus non-metastatic pNETs. The running enrichment score (green line) is plotted as a function of the ranked gene list, with the vertical bars indicating positions of efferocytosis genes and the normalized enrichment score (NES) summarizing pathway-level upregulation in metastatic tumors. **C.** Hallmark "Inflammatory Response" gene set derived from the same integrated analysis. The upper bar plot shows mutation frequencies for selected inflammatory-response genes in metastatic and non-metastatic tumors, as in panel A. The lower heatmap depicts expression profiles of inflammatory-response DEGs across the cohort, using the same color scale. **D.** Hallmark "E2F Targets" gene set analyzed analogously. Mutation frequencies for representative E2F target genes are shown in the upper bar plot, and the lower heatmap displays expression of E2F target DEGs in metastatic and non-metastatic tumors.

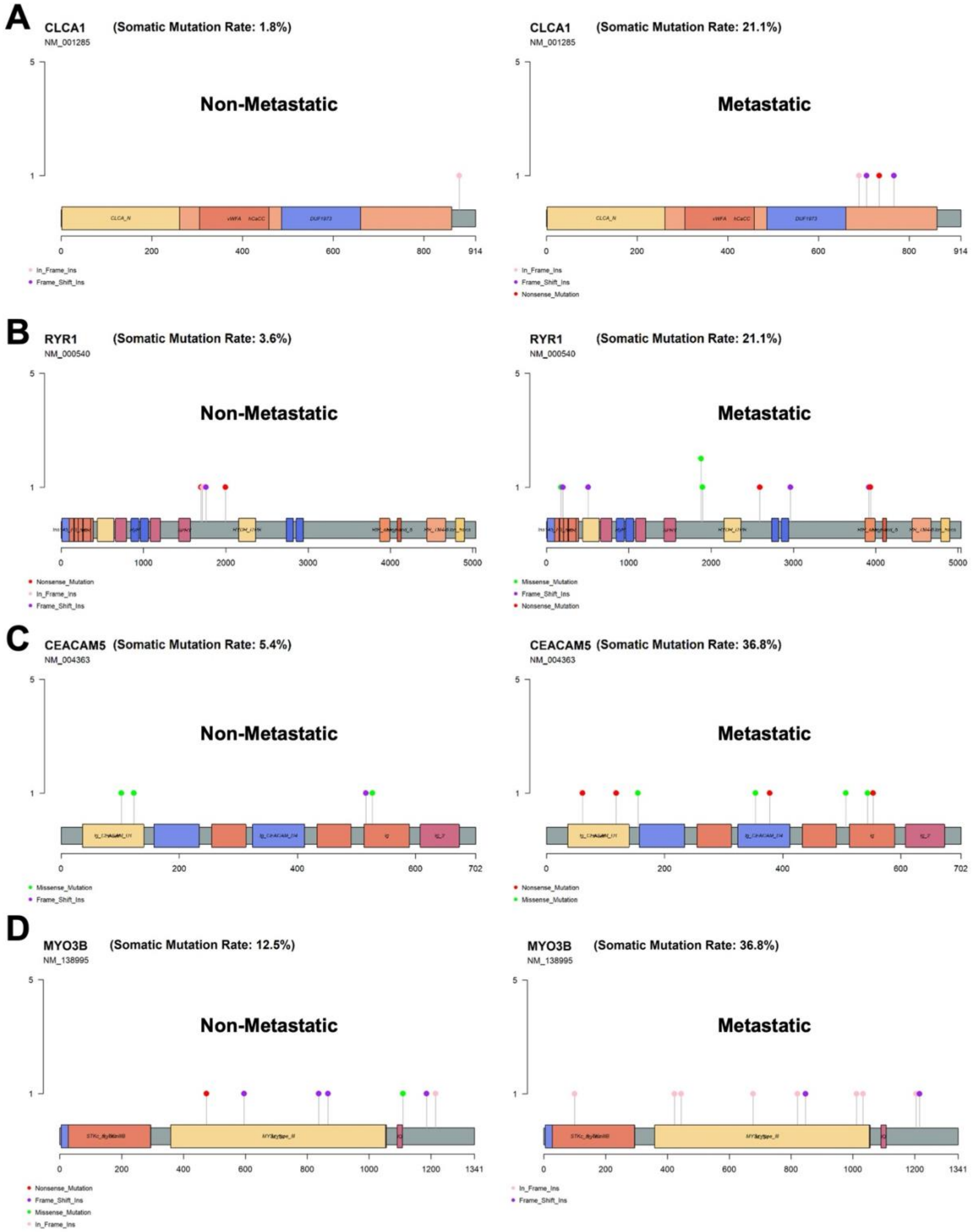


**Supplementary Figure 6. Enrichment and qualitative shift of CLCA1, RYR1, and CEACAM5 coding alterations in metastatic MEN1-wild-type low-grade pNETs.** Lollipop plots depict somatic mutations in selected genes in primary human pNETs profiled by whole-exome sequencing (non-metastatic, n = 56; metastatic, n = 19). For each gene, the left plot shows non-metastatic tumors, and the right plot shows metastatic tumors. The horizontal schematic represents the annotated protein with major domains indicated and amino-acid positions shown on the x-axis; each lollipop marks the position of a distinct variant. Colors denote mutation class as indicated in the below. Somatic mutation rates for each gene and group are reported in the panel titles. **A.** CLCA1 (NM_001285) mutation distribution in non-metastatic (left, somatic mutation rate 1.8%) and metastatic (right, 21.1%) pNETs. The linear map highlights the CLCA_N, vWFA, hCaGC, and DUF1973 domains, showing a single in-frame insertion in non-metastatic tumors versus multiple in-frame and frameshift insertions and nonsense mutations concentrated toward the C-terminal region in metastatic tumors. **B.** RYR1 (NM_000540) mutation distribution in non-metastatic (left, 3.6%) and metastatic (right, 21.1%) pNETs. The full-length ryanodine receptor channel is shown with annotated channel and regulatory domains. Non-metastatic tumors harbor a small number of nonsense, in-frame insertion, and frameshift insertion events restricted between SPRY and RYDR_ITPR domains, whereas metastatic tumors exhibit a higher burden of missense, frameshift insertion, and nonsense mutations distributed heterogeneously across the domains. **C.** CEACAM5 (NM_004363) mutation distribution in non-metastatic (left, 5.4%) and metastatic (right, 36.8%) pNETs. The schematic shows the Ig-like CEACAM domains (e.g., Ig_CEACAM_D1 and Ig_CEACAM_D4) and C-terminal segments. Non-metastatic tumors contain infrequent missense and frameshift insertions, whereas metastatic tumors show a markedly higher frequency of missense and nonsense substitutions along the extracellular domain. **D.** MYO3B (NM_138995) mutation distribution in non-metastatic (left, 12.5%) and metastatic (right, 36.8%) pNETs. The schematic shows mutations in different MYO3B domains and C-terminal segments. Non-metastatic tumors contain frequent frameshift insertions, whereas metastatic tumors shows in frame insertions.

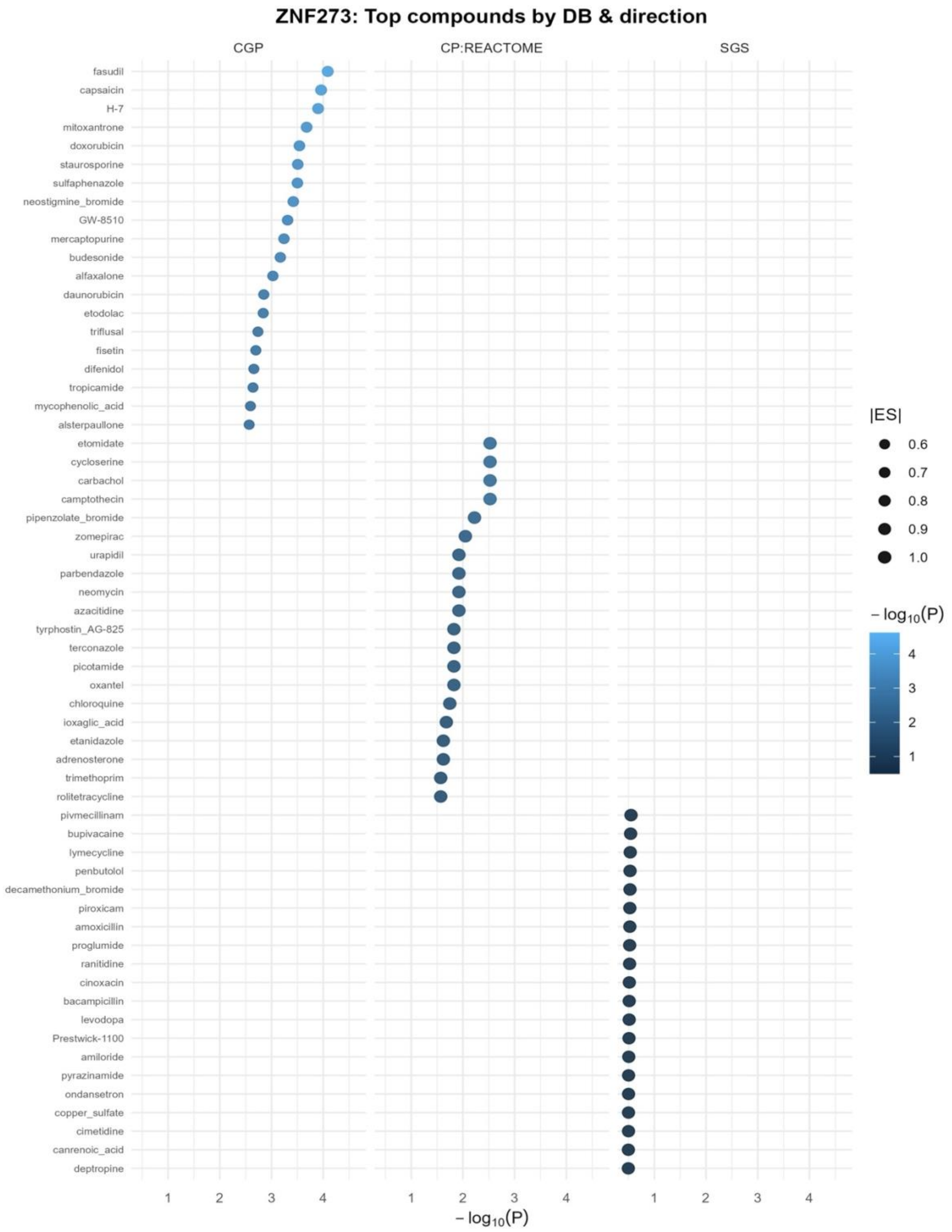


**Supplementary Figure 7. Gene2Drug-predicted compounds targeting ZNF273-associated transcriptional programs.** Bubble plot summarizing small-molecule compounds predicted by Gene2Drug to significantly counteract the gene-expression signature centered on ZNF273 in metastatic pNETs. The y-axis lists individual compounds, and the x-axis shows the statistical significance of their enrichment, expressed as $-\log_{10}(p\text{-value})$. Points are grouped by annotation database used for the enrichment analysis: CGP (cancer gene perturbation signatures), CP:REACTOME (Reactome pathway-based signatures), and SGS (single-gene signatures). Each bubble represents a compound-database pair, bubble size encodes the absolute enrichment score (ES), and color intensity reflects $-\log_{10}(P)$, with darker shades indicating more significant associations.

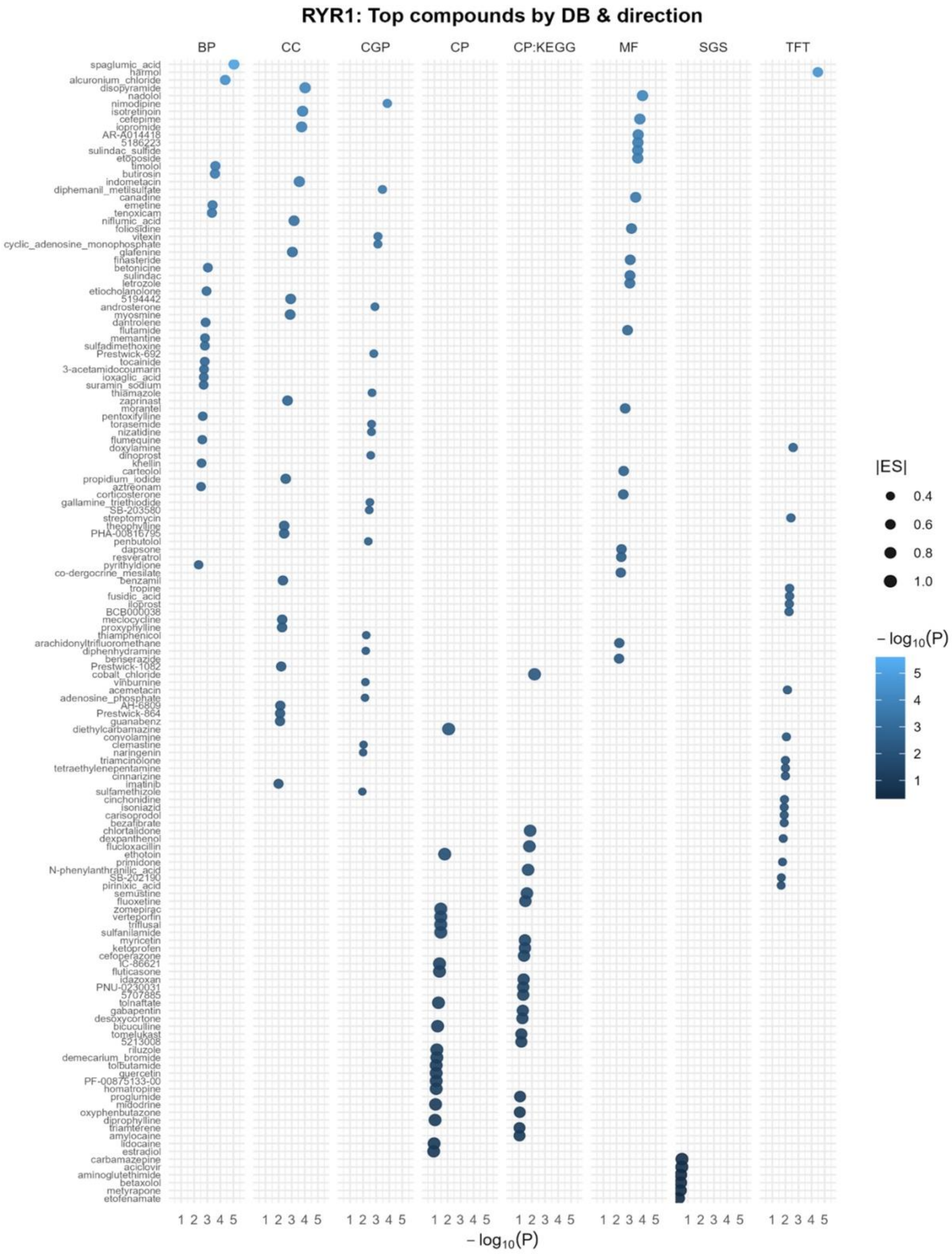


**Supplementary Figure 8. Gene2Drug-predicted compounds targeting RYR1-associated transcriptional programs.** Bubble plot summarizing small-molecule compounds predicted by

Gene2Drug to significantly counteract the RYR1-centered gene-expression signature in metastatic pNETs. The y-axis lists individual compounds, while the x-axis shows $-\log_{10}(p\text{-value})$ for the enrichment of each compound's perturbational profile. Bubbles are grouped into vertical columns according to the annotation database used: Gene Ontology Biological Process (BP), Cellular Component (CC), cancer gene perturbation signatures (CGP), Canonical Pathways (CP), KEGG pathways (CP:KEGG), Molecular Function (MF), single-gene signatures (SGS), and transcription-factor-target sets (TFT). Each bubble represents a compound-database pair; bubble size encodes the absolute enrichment score (ES), and color intensity reflects statistical significance ($-\log_{10}(P)$), with darker shades indicating more significant associations.
